# Retargeted adenoviruses for local IgA and CD47 blocker production as a novel cancer therapy

**DOI:** 10.1101/2025.11.07.686998

**Authors:** M. Chernyavska, K.P. Hartmann, J.H.M. Jansen, N. Baumann, J. Kolibius, D. Brücher, T. Kristoforus, R.H.W. Peters, L. Huijs, D. Laarveld, F. Weiss, R. Burger, M. Lustig, N. Gimenez de Assis, M. Schmid, J.H.W. Leusen, T. Valerius, A. Plückthun, W.P.R. Verdurmen

## Abstract

Despite advances in IgG-based cancer immunotherapy, challenges remain in effectively engaging innate immune responses against solid tumors. Here, IgA antibodies hold promise due to their ability to activate neutrophils and macrophages. We present a novel retargeted adenovirus-mediated approach that transforms cancer cells into “biofactories” for localized production of monomeric or dimeric IgA antibodies and a CD47 blocker to potentiate the effect of IgA antibodies. With our approach tumor cells effectively produced IgA antibodies against tumor antigens such as EGFR or EpCAM and a soluble SIRPα-Fc fusion protein, which blocks the CD47-SIRPα axis. In a perfused tumor-on-a-chip model, locally produced IgA triggered neutrophil- and macrophage-mediated tumor cell killing, further potentiated by SIRPα-Fc co-production. In FcαRI-transgenic, tumor-bearing mice, intratumoral adenoviral injection induced strong local IgA and SIRPα-Fc expression, immune cell infiltration, and more than 50% tumor volume reduction after a single treatment. We found that dimeric IgA exerts stronger effects than monomeric IgA, which is of particular interest since dimeric IgA necessitates a local production approach. Together, these results demonstrate that adenovirus-mediated, tumor-restricted delivery of IgA antibodies and CD47 blockade effectively engages innate immune mechanisms, providing a promising new avenue to enhance cancer immunotherapy.

**Graphical abstract:** 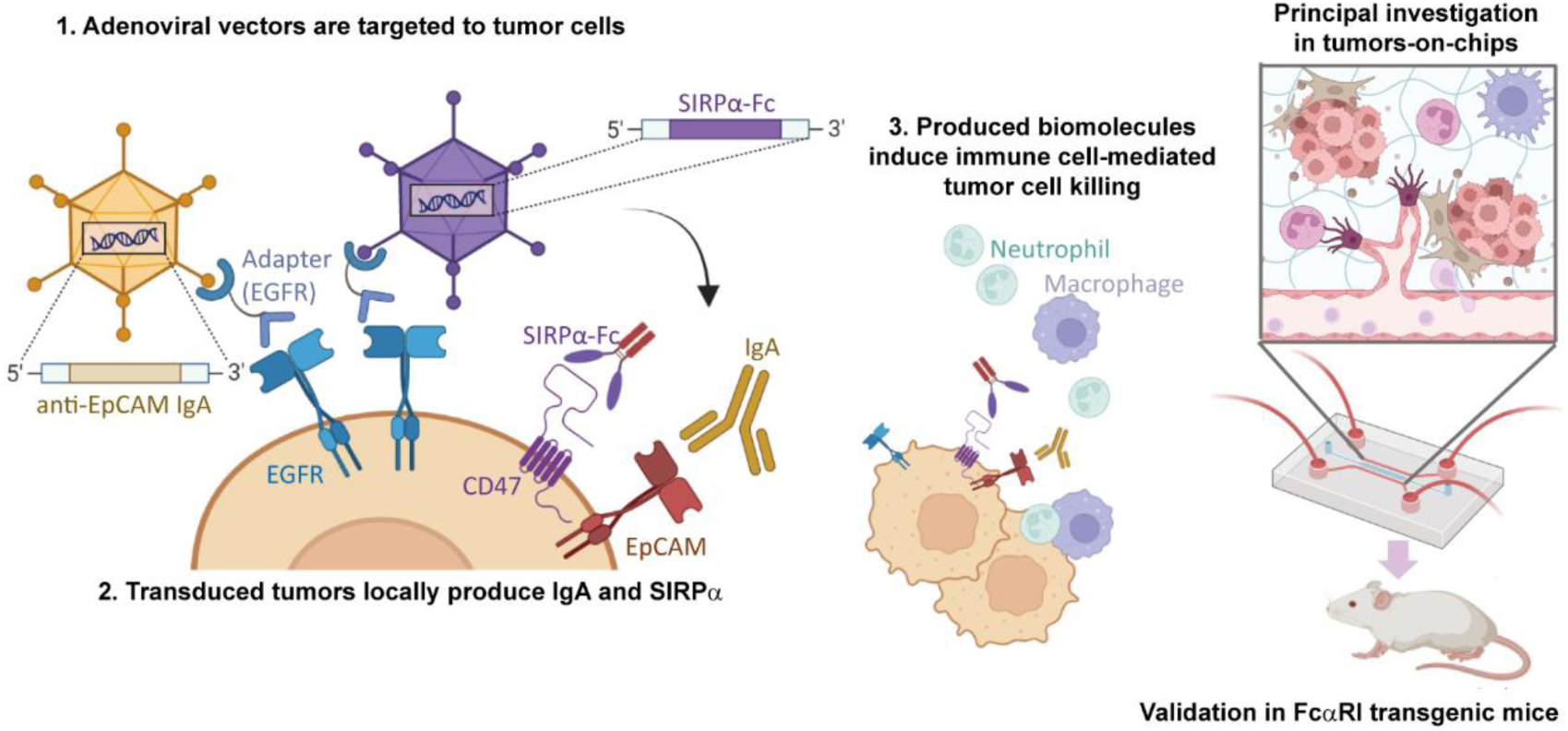

## 1. Introduction

Cancer immunotherapy has emerged as a powerful treatment approach, with current clinical antibodies primarily based on the IgG isotype.^1,2^ However, also IgA antibodies have shown potential to fight cancer by effectively activating myeloid cells — key components of the innate immune system — with particularly improved activation of neutrophils compared to IgG molecules.^3–5^ While IgG antibodies bind to a widely expressed and heterogeneous family of Fcγ receptors,^6,7^ which primarily activate macrophages and natural killer (NK) cells for tumor cell killing, monomeric and dimeric IgA antibodies engage a single Fcα-receptor (FcαRI; CD89), which is almost exclusively expressed by myeloid cells, and its engagement potently activates neutrophils and macrophages.^8^

Neutrophils, the most abundant immune cells in human circulation, play a complex role in cancer.^9–11^ While they can effectively kill cancer cells,^12,13^ they are often associated with poor prognosis due to their potential to promote tumor growth and spread, caused by their secreted cytokines and ability to release neutrophil extracellular traps (NETs).^13,14^ Activating neutrophils to attack tumors, rather than sustain tumor growth, represents a promising strategy for IgA-based therapies.^2,4,15^ Studies have shown that IgA-activated neutrophils can eliminate cancer cells through trogoptosis (killing by trogocytosis) – a process of “frustrated phagocytosis”, by which neutrophils nibble off parts of cancer cell membranes.^16^ Similarly, macrophages frequently promote tumor growth within the tumor microenvironment by secreting cytokines that stimulate tumor progression and immune evasion. However, they can also be activated to eliminate cancer cells through antibody-dependent cell-mediated phagocytosis (ADCP).^17–19^ ADCP has been observed to be mediated by both IgG and IgA antibodies to a similar level.^20–23^ Notably, IgA has demonstrated superior ability to activate tumor cell killing compared to IgG in whole blood assays,^11,24,25^ and also showed efficacy *in vivo.*^26–28^ Furthermore, dimeric IgA, which is the most common form in mucosal tissues, is even more effective than monomeric IgA in Fc and F(ab)-mediated tumor cell killing mechanisms, the latter referring to mechanisms involving, for instance, inhibition of ligand binding and receptor downmodulation.^22^ To date, monomeric IgA efficacy for elimination of cancer cells has been shown for a number of targets, including epidermal growth factor receptor (EGFR), human epidermal growth factor receptor 2 (HER2), the B-cell antigen CD20 and ganglioside GD2, both *in vitro* and *in vivo.*^3,11,27–31^

The anti-cancer effects of IgA can be further enhanced by blocking the CD47-signal-regulatory protein alpha (SIRPα) innate immune checkpoint.^28,32,33^ CD47 on tumor cells binds to SIRPα (CD172A) on myeloid effector cells and triggers ITIM-dependent signaling cascades, thereby, acting as a “don’t eat me” signal on cancer cells, helping them to evade destruction by the immune system. Blocking this signal with different inhibitors can boost both innate and also adaptive immune responses,^34^ and has shown promise in many preclinical and early clinical trials.^35^ This has led to the development of various CD47 and SIRPα inhibitors for use in combination with IgG-based immunotherapies,^32,36,37^ but these molecules are also highly effective in the context of IgA antibodies.^28,38,39^

IgA therapeutics are limited clinically by a short plasma half-life^26^ and the lack of FcαRI in wild-type mice,^40^ complicating preclinical studies as transgenic FcαRI-expressing mice are typically required.^41,42^ Due to the ubiquitous expression of CD47, its blockade has been linked to systemic side effects, leading particularly to anemia and thrombocytopenia.^35^ To address these challenges, we developed a novel approach that focuses on producing IgA antibodies and a CD47 blocker directly within the tumor microenvironment. This localized production is achieved using a modified human adenovirus of serotype 5 with a previously developed adapter technology, to direct the adenovirus to specifically bind to cancer cells in a receptor-specific manner, while minimizing liver transduction and interactions with immune components.^43,44^ This approach was successfully used for delivery and in situ-production of chemokines, cytokines and of an anti-HER2 IgG gene and showed therapeutic efficacy *in vivo,*^45,46^ as well as for cell-specific gene delivery to T cells and dendritic cells.^47,48^

In our study, we adapted this technology to produce both monomeric and J-chain-containing dimeric forms of IgA as well as a CD47 blocker in the tumor. To gain a deeper understanding of this therapeutic approach, we first utilized a perfusable tumor microenvironment-mimicking tumor-on-a-chip model before moving to *in vivo* studies.^49,50^. We chose to target EGFR and EpCAM using both monomeric and dimeric forms of IgA antibodies, in combination with simultaneous production of a soluble SIRPα-Fc protein, which blocks the CD47-SIRPα axis. Using our tumor-on-a-chip model, we gained detailed insights into the mechanisms behind this enhanced efficacy. Finally, we validated our findings *in vivo* in mice, demonstrating successful production of IgA antibodies and the SIRPα-Fc protein within xenografted tumors and robust therapeutic activity after only a single injection.

## 2. Results

### 2.1. Vector engineering for production of diverse IgA antibody formats and a CD47 blocker

To produce the IgA antibodies and CD47 blocker for our study, we utilized a multicistronic vector system. This system allows for efficient production of both the heavy and light chains of an antibody from a single genetic construct, ensuring optimal antibody assembly and function.^51^ We first adapted this system to produce an anti-EGFR IgA antibody with an optimized framework (IgA2.0) previously shown to enhance IgA stability and *in vivo* half-life.^5^ Both HC and LC were expressed under a single promoter, separated with a 2A “self-cleaving” (ribosome-skipping) peptide sequence (Figure S1). Next, we used this system to engineer a novel IgA against human EpCAM in the IgA2.0 framework using the variable domains from a previously reported single-chain variable fragment against human EpCAM,^52^ constructing again an IgA monomer as in Figure S1a. The original single-chain variable fragment has previously been extensively characterized and was found to have low-nanomolar affinity and high specificity, both *in vitro* and *in vivo.*^52,53^ Both anti-EGFR and anti-EpCAM IgA antibodies were successfully produced in CHO-S cells upon plasmid transfection, and their binding activity and specificity were confirmed by ELISA (Figure 1a). Upon comparing P2A and T2A “self-cleaving” peptide sequences, we observed higher expression of IgA from P2A-variant constructs, and these were selected for further studies (Figure S2). Both antibodies showed the expected patterns in western blots (Figure 1b).

**Figure 1.**
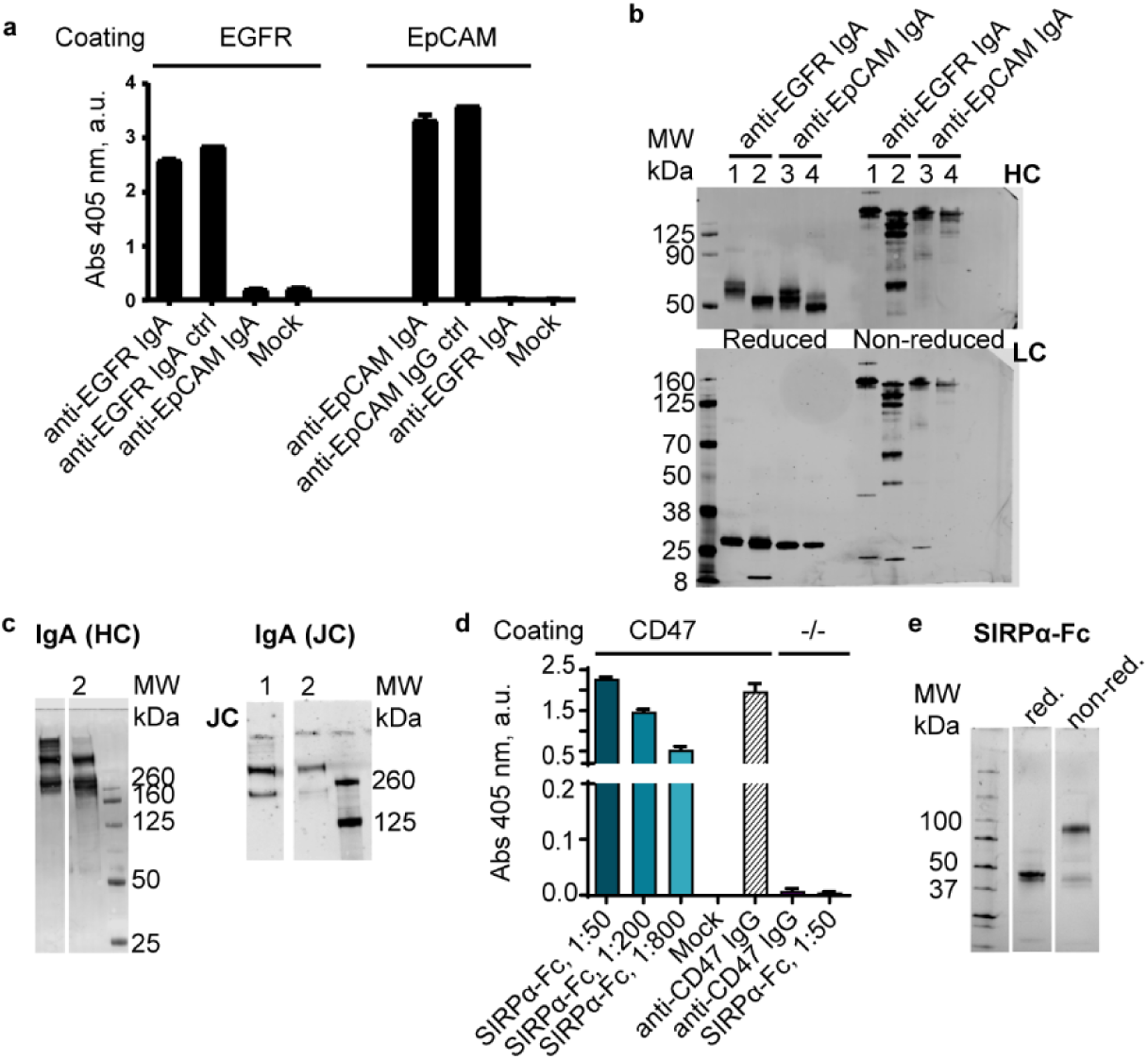
Characterization of engineered IgA antibodies and the CD47-blocking SIRPα-Fc protein. **(a)** ELISA was used to measure the target-binding activity of anti-EGFR and anti-EpCAM IgA antibodies derived from the supernatants of transfected CHO-S cells. Recombinant IgA and IgG antibodies were used as positive controls (ctrl: 0.1 nM), and supernatants from mock-transfected cells served as negative controls. Binding specificity was confirmed by assessing the binding of each IgA antibody to its intended target (EGFR or EpCAM) and to the non-target protein. **(b)** Western blot analysis of anti-EGFR and anti-EpCAM IgA antibodies in the supernatant of transfected CHO-S cells (lane 1,3). Purified recombinant IgA antibodies were used as controls (lane 2,4). Note that the anti-EGFR IgA control shows incomplete antibody assembly. HC, heavy chain; LC, light chain. **(c)** Western blot analysis of dimeric IgA antibodies under non-reducing conditions, showing the presence of both the IgA heavy chain (HC), detected by a mouse anti-human kappa chain antibody, and the J-chain (JC), detected by a mouse anti-poly His-tag antibody, in supernatants of CHO-S transiently transfected with the respective expression vectors. 1: anti-EGFR; 2: anti-EpCAM. **(d)** ELISA was performed to measure SIRPα-Fc protein levels in the supernatant of transfected CHO-S cells, using CD47 as coating. A recombinant anti-CD47 IgG antibody (2.5 nM) was used as a positive control, and supernatant from mock-transfected cells served as a negative control. Wells with no CD47 coating served as additional controls. **(e)** Western blot analysis of the SIRPα-Fc protein, as detected by a mouse anti-γ heavy chain, in supernatants of CHO-S cells transiently transfected with the respective expression vector, under reducing (red.) and non-reducing (non-red.) conditions. (a-e): representative images from at least three independent experiments are shown.

Next, we engineered a dimeric format of each of the antibodies by introducing a joining chain (J-chain) downstream of the IgA expression cassette following an EMCV IRES sequence (Figure S1). The dimeric antibodies were successfully produced, as evidenced by their molecular weights seen in western blotting (Figure 1c). For the dimeric construct, the T2A sequence was selected due to better production with this sequence (Figure S3). Interestingly, we observed some potential aggregation with the anti-EGFR IgA dimer, which was less prominent for the anti-EpCAM IgA dimer. Notably, the dimeric fraction was between 30-50% of the total detected IgA protein. Target binding of monomeric and dimeric IgA on cells was confirmed through immunofluorescence using A-431 and MCF-7 cells, overexpressing EGFR and EpCAM (Figure S4).

Finally, as a CD47 blocker, we encoded a previously reported SIRPα-Fc fusion molecule which is also cross-reactive with murine CD47, albeit with a somewhat reduced affinity (1 nM for human CD47 vs. 58 nM for murine CD47)^34^ (Figure S1). This protein was designed with an FcRn-binding inactive Fc region (AAA-N297A mutant Fc of the IgG1^54^), to avoid unwanted side effects due to the ubiquitous presence of CD47 on healthy cells, while being stable in the plasma.^36^ The proper production of the SIRPα-Fc fusion protein was confirmed by analysis of its CD47-binding activity by ELISA (Figure 1d), its molecular weight by western blotting (Figure 1e), and its cross-reactivity with murine CD47 by immunofluorescence using murine MC-38 cells, with OVCAR3 acting as a high CD47-expressing human reference cell line (Figure S4).

After having demonstrated the functionality of our proteins through transient transfections, we affinity-purified the IgAs by protein L affinity chromatography and the CD47 blocker by protein A chromatography. The Fab and Fc functions of IgA were further characterized with ^51^Cr release-based ADCC assays with human neutrophils on MDA-MB-468 and HCT116 cell lines, which we selected because they express high levels of both EGFR and EpCAM,^55–58^, the targets for our IgA antibodies, as well as CD47, the target for our CD47 blocker.^59,60^ The ADCC assay confirmed the ability of the engineered IgA antibodies to induce neutrophil-mediated ADCC of tumor cells (Figure S5), with the exception of the purified dimeric mix of anti-EGFR IgA for which we observed aggregation. We then proceeded to generate the adenoviral (AdV) vectors encoding the proteins using the modified pAdEasy-1 HRV adenoviral genome, with purified proteins serving as benchmarks for comparing the activity of AdV vector-mediated locally produced proteins. In total, five AdV vectors were made encoding (i) monomeric anti-EGFR IgA, (ii) monomeric anti-EpCAM IgA, (iii) dimeric anti-EGFR IgA, (iv) dimeric anti-EpCAM IgA and (v) SIRPα-Fc, using the engineered pShuttle expression vectors (Figure S1).

### 2.2. Adenovirus-mediated delivery of therapeutic genes in the tumor microenvironment

To validate the AdV-mediated IgA gene delivery approach, the AdV vectors were retargeted to either EGFR or EpCAM, using the molecular adapter retargeting platform reported earlier^43,44^ (Figure 2a). To provide proof of concept for the IgA-based therapy using the AdV platform, we again employed MDA-MB-468 and HCT116 cells.

**Figure 2.**
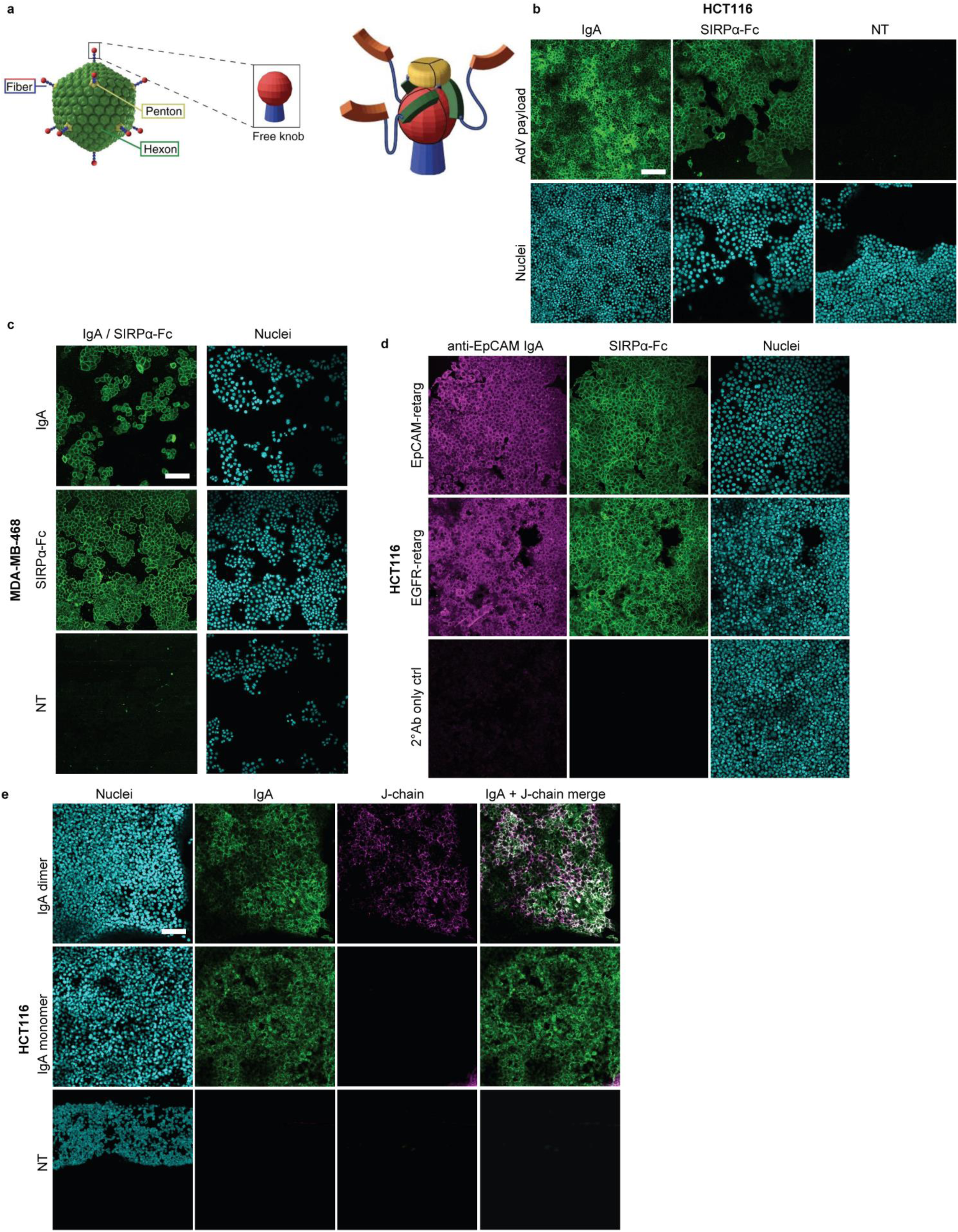
Production and detection of monomeric and dimeric IgA antibodies, as well as SIRPα-Fc protein upon retargeted AdV transduction. **(a)** Schematic image showing the retargeting approach of the adenovirus. The adenovirus capsid is composed of hexons forming its main shell, pentons anchoring at the vertices, and fibers projecting from those vertices for host cell attachment. As reported in previous work, a trimeric DARPin (orange) can be engineered to cover the free knob at the tip of each fiber of the adenovirus in order to achieve retargeting^43^. **(b,c)** Monomeric anti-EpCAM IgA and anti-EGFR IgA were produced in HCT116 **(b)** and MDA-MB-468 cells **(c)**, respectively, via retargeting of the AdV vector to EGFR or EpCAM using the DARPin-based approach described in **(a).** Green, AdV payload (IgA, labeled by anti-kappa-light chain, or SIRPα-Fc, labeled by anti-human IgG. Secondary antibodies were Alexa-488 labeled); cyan, nuclei (Hoechst). MOI: 50 i.v.p./cell (IgA), 5 i.v.p./cell (SIRPα-Fc). **(d)** Comparison of adenovirus-mediated co-delivery of IgA and SIRPα-Fc when retargeting the respective adenovirus to EpCAM or EGFR (EpCAM- or EGFR-retarg) on the surface of the cancer cells (HCT116), which express both receptors**. (e)** Anti-EpCAM IgA dimer was delivered via EpCAM-retargeted AdV in HCT116 cells. J-chain (magenta) was detected in the dimer-transduced sample, and co-localized with IgA (green) signal. Cyan, nuclei (Hoechst). Positive controls (recombinant IgA, SIRPα-Fc proteins) and secondary Ab only control images are shown in Figure S6. All cells are non-permeabilized, meaning products we see have been secreted from the cells and re-bind from the outside in an autocrine or paracrine fashion. Representative images from two independent experiments are shown. NT, non treated. Scale bars: 100 μm.

#### 2.2.1. Autocrine delivery in 2D

We first confirmed that our modified AdV vectors could effectively deliver the antibody genes and produce IgA antibodies within the tumor cells. After addition of the AdV vectors encoding anti-EpCAM or anti-EGFR IgA to HCT116 and MDA-MB-468 cells, retargeted to EGFR and EpCAM, respectively, we observed strong binding of the in situ-produced IgA antibodies to the surface of non-permeabilized cells, confirming their expected extracellular localization (Figure 2b,c; Figure S6). To examine the effect of receptor targeting, we delivered anti-EpCAM IgA to HCT116 cells using AdV vectors retargeted to either EpCAM or EGFR. Both approaches resulted in comparable levels of IgA bound to its target (Figure 2d). Therefore, to avoid receptor competition in subsequent experiments, we chose to target the viral vector to the alternative receptor—e.g., delivering anti-EGFR IgA via targeting to EpCAM, and vice versa. Next, we confirmed the production of dimeric IgA by staining simultaneously for IgA and the J-chain (Figure 2e). Finally, we verified that delivery of the AdV vector encoding the SIRPα-Fc gene led to successful production of the SIRPα-Fc protein, which bound CD47 on both cell lines (Figure 2b,c; Figure S6).

#### 2.2.2. Paracrine (co-) delivery

We next investigated whether the therapeutic proteins produced upon transduction with the AdV vector could spread to neighboring tumor cells, creating a “bystander effect”. To test this, we introduced the AdV vector into a population of labeled tumor cells and then added a second population of unlabeled, non-transduced cells. We found that the IgA antibodies and SIRPα-Fc protein produced by the labeled cells were also present on the surface of the non-transduced (unlabeled) cells (Figure S7-8), demonstrating that these therapeutic proteins can effectively diffuse towards neighboring tumor cells. We then explored the possibility of simultaneously delivering both the IgA antibody and the CD47 blocker using a combination of two AdV vectors. This approach resulted in successful co-expression and distribution of both therapeutic proteins on the transduced tumor cells and their non-transduced neighbors (Figure S7-8).

#### 2.2.3. AdV-mediated IgA and CD47 production in a tumor-on-a-chip model

To further evaluate our adenoviral vector delivery system in a setting closer resembling *in vivo* physiology, we used a tumor-on-a-chip model. This model mimics the tumor microenvironment more accurately than traditional cell culture methods, as we can incorporate non-cancerous cells, extracellular matrix components and fluid flow, which are factors impacting therapeutic delivery and efficacy.^61,62^ Our in-house designed tumor-on-a-chip model features tumor cells (MDA-MB-468 or HCT116) and non-malignant cells of the tumor microenvironment (endothelial cells [HUVEC] and fibroblasts [C5120] combined in an extracellular matrix [ECM, collagen] in the tissue compartment), recapitulating a human-like tumor microenvironment. The media is provided from two side channels connected by microtunnels to the tissue compartment.^63^ The media flow is induced by bidirectional rocking of the device facilitating the nutrient supply, exchange, as well as improving diffusion of the compounds into the tissue.^64^ Using this model, we confirmed that the AdV vector could effectively deliver the genes and yield production of both the IgA antibodies and the SIRPα-Fc fusion protein within this more complex environment (Figure 3a,b). Importantly, the IgA antibodies specifically targeted the cancer cells and did not bind to the non-cancerous cells, while the SIRPα-Fc protein, as expected, bound to all cell types due to the widespread presence of CD47 (Figure 3b, Figure S9). We also confirmed successful co-delivery of both therapeutic proteins in this model as illustrated by a representative slice (Figure 3c, see Figure S9 for additional controls) and an orthogonal section and 3D reconstruction of the tumor-on-a-chip model (Figure 3d-f).

**Figure 3.**
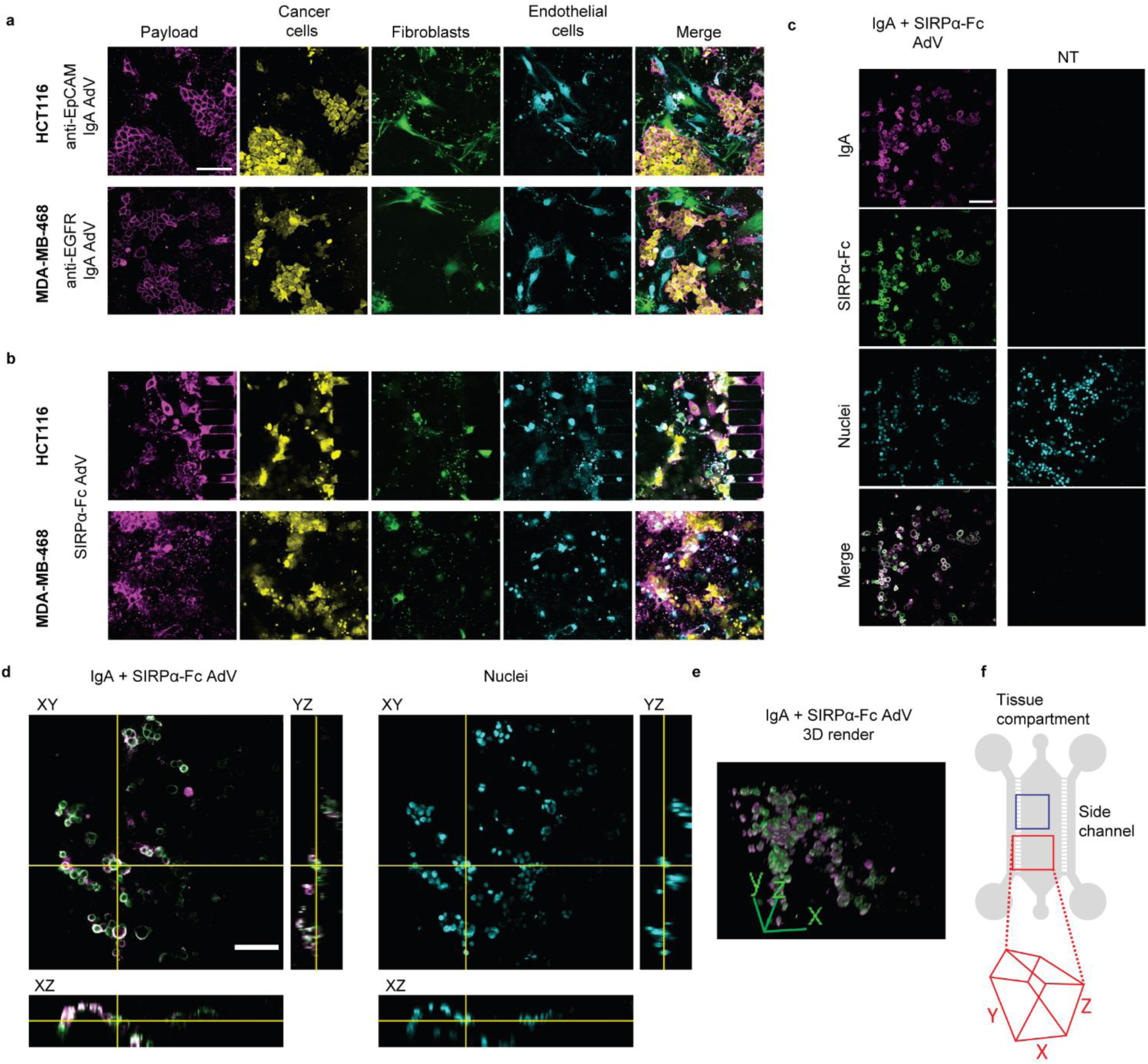
Adenovirus-mediated local IgA and SIRPα-Fc production and binding specificity was analyzed in a tumor microenvironment (TME) on-chip. Cells were stained with different CellTrace dyes, proteins were detected with fluorescently labeled antibodies. **(a)** Anti-EpCAM IgA or anti-EGFR IgA (Payload, magenta) was locally produced in the TME featuring MDA-MB-468 or HCT116 cells (Cancer cells, yellow), respectively, as well as fibroblasts (green) and endothelial cells (cyan). IgA-encoding adenoviruses were retargeted with the respective adapters to EGFR or EpCAM expressed on HCT116 or MDA-MB-468 cells, respectively, at an MOI of 20 i.v.p./cell. **(b)** Similar to (a), SIRPα-Fc was produced *in situ* upon AdV retargeting to EGFR (HCT116) or EpCAM (MDA-MB-468) at an MOI of 20 i.v.p./cell. The overlay image shows no specificity of SIRPα-Fc towards either of the cell types, due to ubiquitous expression of CD47 on all cells. **(c)** Anti-EGFR IgA (magenta) and SIRPα-Fc (green) simultaneously produced *in situ* upon co-delivery with EpCAM-retargeted adenoviral vectors in MDA-MB-468 (nuclei: cyan, Hoechst) at an MOI of 20 i.v.p./cell (each). Merge depicts SIRPα-Fc and IgA. **(d)** Adenovirus-mediated co-delivery of IgA and SIRPα-Fc as in (c), represented as orthogonal sections showing 3D distribution of cells with the *in situ* produced payloads on the cell surface, with the 3D reconstruction shown in **(e)** (dimensions indicated). Positive controls (recombinant IgA, SIRPα-Fc) and secondary Ab only control images are shown in Figure S9. **(f)** A schematic image of the previously utilized tumor-on-a-chip (not drawn to scale), consisting of a main chamber with cell co-culture in collagen, and two side channels with medium reservoirs through which retargeted adenoviral vectors or control proteins are administered^63^. Representative images are shown, n=4. Scale bars: 100 μm.

### 2.3. AdV-based local IgA antibody delivery induces tumor cell killing by human neutrophils on-chip

To assess the ability of our AdV-based local IgA antibody production to activate neutrophils to kill cancer cells, we developed an image-based assay within our tumor-on-a-chip model (Figure 4a-d, Figure S10). This assay mimics an important element of the natural tumor immune microenvironment through the incorporation of neutrophils, which can enter and migrate through the tissue and interact with cancer cells. We used a physiologically relevant concentration of neutrophils (10 million/mL), comparable to what is found in the bloodstream (1.5–8.5 million/mL).^65^

**Figure 4.**
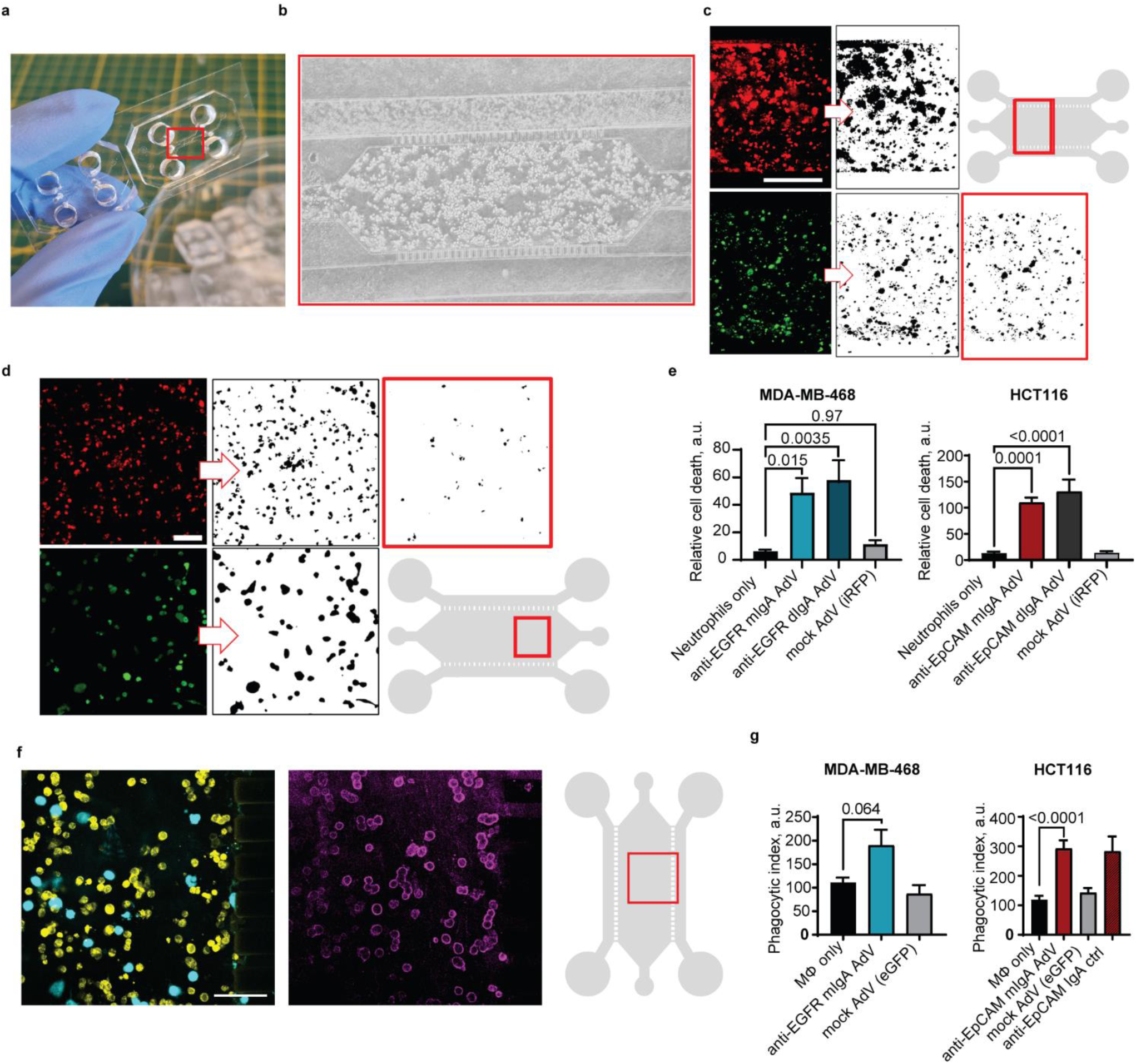
IgA antibodies produced by AdV-transduced cancer cells activate immune cells to kill cancer cells in tumor-on-a-chip models via ADCC and ADCP. **(a)** A photograph of the polydimethylsiloxane (PDMS) microfluidic device with the inset imaged microscopically in **(b)**. The microphotograph shows the tissue compartment with tumor cells embedded in collagen, and two side perfusion channels. Primary human neutrophils were perfused through the upper side channel. **(c)** Method for visualization of antibody-dependent cell-mediated cytotoxicity (ADCC). Cells were stained with different Cell Trace dyes. Here, tumor cells are labeled red, and dead cells are labeled green (CellTox Green). Black and white images represent cell masks. The red outline highlights dead tumor cells, being recognized as the intersection from the green and red channel signals. Scale bar: 400 μm. The schematic of the chip shows an approximate area visualized (red inset). **(d)** Method of antibody-dependent cellular phagocytosis (ADCP) on-chip visualization and processing based on the labeling of the tumor cells (red) and macrophages (green). Schematic of the chip as in (c); scale bar: 100 µm. **(e)** Quantification of ADCC activity by microscopy-based ADCC assay (see Methods). The graph shows the ability of neutrophils to kill MDA-MB-468 and HCT116 tumor cells after the tumor cells are transduced with AdV vectors encoding genes for anti-EGFR or anti-EpCAM IgA antibodies (MOI 50 i.v.p./cell). Controls include tumor cells with neutrophils only and tumor cells transduced with an irrelevant protein (mock AdV). "mIgA" and "dIgA" refer to monomeric and dimeric IgA, respectively. Error bars represent standard error of the mean (SEM). Data are from at least 5 experiments using cells from 3 different blood donors. Statistical analysis was performed using one-way ANOVA with Dunnett’s correction for multiple comparisons. Adjusted p-values are shown. **(f)** Confocal microscopy images of HCT116 tumor cells (yellow) and macrophages (cyan) within the tumor-on-a-chip model. The left image shows the cells labeled with cytoplasmic dyes. The right image shows staining for anti-EpCAM IgA antibodies (magenta) produced by the tumor cells 24 h after transduction with the adenoviral vector (MOI 50 i.v.p./cell). Scale bar: 100 μm. **(g)** Quantification of ADCP activity. The phagocytic index (see Methods) of macrophages (MФ), expressed in arbitrary units (a.u.), is shown for MDA-MB-468 and HCT116 models upon treatment with anti-EGFR IgA or anti-EpCAM IgA, respectively, with genes delivered via retargeted adenovirus. Controls include untreated tumor cells, tumor cells transduced with a mock AdV, and tumor cells treated with purified anti-EpCAM IgA antibodies (5 nM). Error bars represent SEM. Data are from at least 5 experiments using macrophages derived from 3 different batches of pooled donors. Statistical analysis was performed using an unpaired t-test. Adjusted p-values are shown.

Initially, we confirmed that our purified IgA antibodies could effectively induce neutrophil-mediated killing of cancer cells in this model (Figure S10). Within hours after the start of the assay with purified anti-EpCAM IgA and anti-EGFR IgA as benchmark proteins added to the side channels, neutrophils were observed to enter the tumor compartment, migrate through the ECM and kill tumor cells (Figure 4, Figure S10). We then tested the ability of the AdV-encoded locally produced IgA antibodies to activate neutrophils. As expected, both anti-EGFR and anti-EpCAM antibodies produced by the AdV vector effectively triggered neutrophil-mediated killing of the respective cancer cells (Figure 4e, Figure S11), as measured by our microscopy-based ADCC assay. In HCT116 cells, anti-EpCAM IgA resulted in an average of 12.6 ± 7.1-fold increase in ADCC over background (neutrophils with cancer cells without treatment) for the monomeric, and 14.2 ± 8.7-fold increase for the dimeric IgA form delivered with EGFR-targeted AdV. Similarly, in MDA-MB-468 cells, anti-EGFR IgA monomer delivered with EpCAM-targeted AdV resulted in relative ADCC on average 6.8 ± 1.8-fold over background; dimeric IgA in the same context resulted in ADCC on average 8.0 ± 2.9-fold higher than background. Importantly, the AdV vector itself did not activate neutrophils, indicating that the observed killing was specifically due to the IgA antibodies. We also observed that targeting either EGFR or EpCAM with the AdV vector resulted in similar levels of neutrophil activation and cancer cell killing (Figure S11).

Furthermore, we analyzed neutrophil-mediated ADCC on-chip with confocal microscopy. The DNA-binding dye CellTox Green, which we initially employed to detect dead cells, revealed the presence of fibrous DNA structures following the direction of neutrophil tissue infiltration and migration through the gel in the tissue compartment, which can be attributed to neutrophil extracellular traps (NETs)^66^ (Figure S12). NETs were observed in the chips with the highest neutrophil ADCC activity and were not found in the chips with no treatment. The NETs were detected “wrapping” the clusters of tumor cells, suggesting NETosis to be one of the mechanistic aspects of neutrophil activation upon IgA treatment, as seen before.^67^

### 2.4. M2-polarized macrophages phagocytose cancer cells in response to AdV-based local IgA delivery

Having demonstrated that our AdV-mediated IgA production approach can activate neutrophils to kill cancer cells, we next investigated their effects on macrophages. Macrophages are another type of immune cell capable of destroying cancer cells through ADCP, particularly anti-inflammatory, M2-like (tumor promoting) macrophages, which we showed to be potently activated by IgA antibodies.^33^ To study this, we used our tumor-on-a-chip model, where cancer cells and macrophages are co-cultured in a more physiological microenvironment. We delivered IgA genes via AdV-mediated transduction of tumor cells in tumor-macrophage co-cultures on-chip. We assessed the ability of macrophages to engulf and kill cancer cells using a confocal imaging-based assay. We confirmed production of anti-EpCAM IgA expression at the moment of the assay (Figure 4f) and found that both anti-EGFR and anti-EpCAM IgA antibodies, produced upon adenoviral transduction, significantly enhanced macrophage-mediated ADCP of MDA-MB-468 (2.1 ± 0.5-fold) and HCT116 cancer cells (2.4 ± 0.6-fold), respectively, compared to non-treated controls (Figure 4g). Recombinant anti-EpCAM IgA alone (in the HCT116 model) resulted in a 2.5 ± 0.3-fold increase in tumor cell phagocytosis over the non-treated control, which is on the level of AdV-mediated *in situ* produced IgA, and is in line with our previously observed phagocytic activity upon anti-EGFR IgA treatment.^33^ The effects can be attributed to the produced antibody and not the viral vector itself, as the mock transduction with AdV not encoding an antibody did not enhance ADCP.

### 2.5. AdV co-delivery demonstrates the therapeutic potential of local CD47 blockade in combination with IgA treatment to enhance ADCC and ADCP of tumor cells

We then established the flexibility of our approach for co-delivery of different AdV payloads to enhance IgA-mediated tumor cell elimination by neutrophils and macrophages. We aimed to illustrate how the previously reported augmentation of IgA-mediated ADCC or ADCP^16,33,37^ by CD47 blockade can also be achieved by co-delivery of IgA and soluble SIRPα-Fc by tumor-targeted AdV vectors in the tumor-on-a-chip model.

First, we tested the effect of SIRPα-Fc co-delivery in ADCC assays in EGFR-positive tumor-on-a-chip models: MDA-MB-468 and A-431 cells, both previously used in demonstrating the effect of CD47 blockade on ADCC.^33^ In both models, when SIRPα-Fc was co-delivered with IgA antibody genes upon AdV transduction, ADCC was significantly enhanced compared to IgA delivery alone (Figure 5a,b). For the co-delivery of SIRPα-Fc and anti-EGFR IgA as purified proteins in the A-431 model, similar effects were seen yet without reaching significance (Figure S13). Comparable effects were observed for dimeric IgA delivery, although significance was only reached for A-431 cells (Figure 5a,b). As expected, AdV-mediated delivery of just CD47 blocker alone, or its administration as a recombinant protein, did not induce ADCC owing to the lack of its FcR-binding function, which has been removed from the SIRPα-Fc fusion protein (Figure 5a,b, Figure S13).

**Figure 5.**
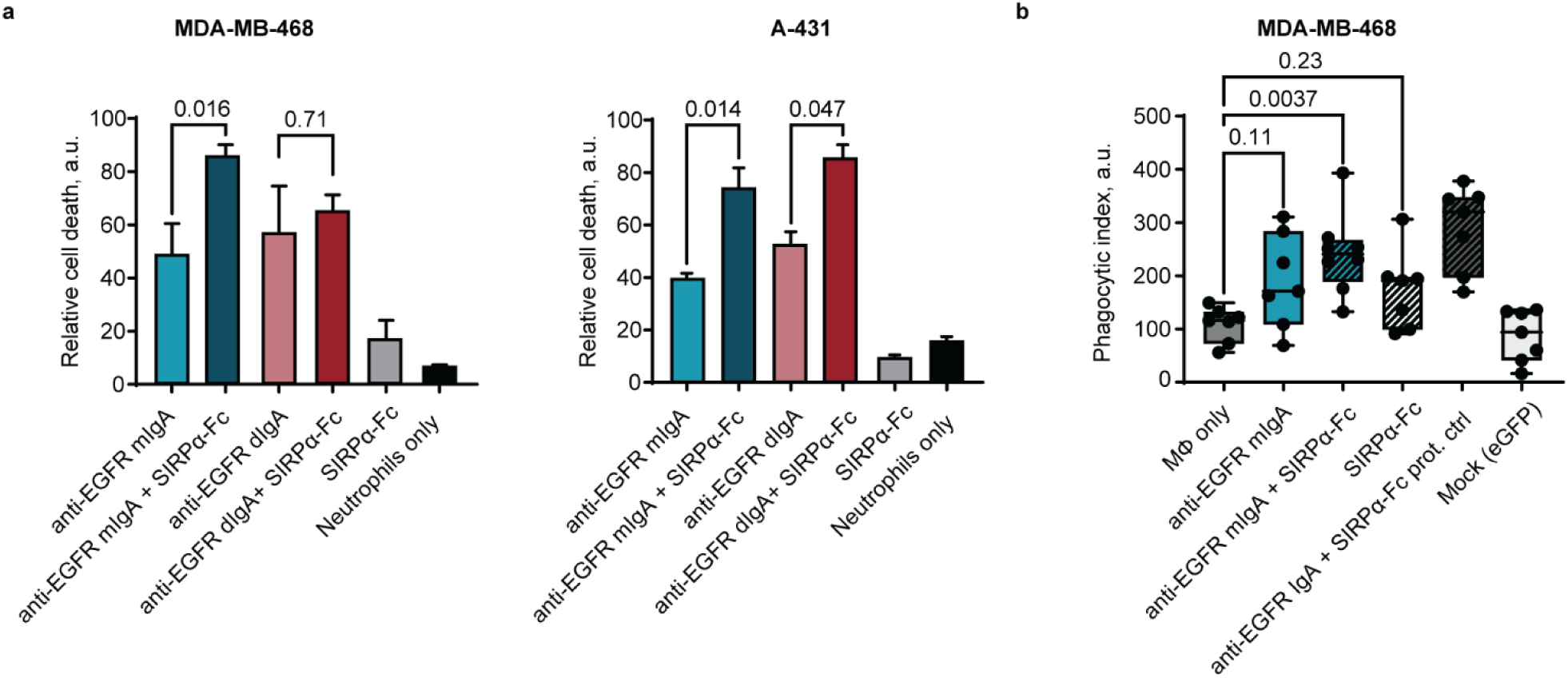
AdV-mediated co-delivery of IgA antibody genes and SIRPα-Fc shows innate immune checkpoint inhibition and enhances ADCC and ADCP. **(a)** Results of the ADCC assay on-chip showing the activity of primary human neutrophils in killing MDA-MB-468 and A-431 cells via AdV-delivered, *in situ* produced anti-EGFR IgA (monomer, mIgA or dimer, dIgA), compared to the combination with the SIRPα-Fc co-delivered with the respective IgA. Administration of the SIRPα-Fc alone was tested as a control condition, and non-treated cancer cells with neutrophils alone were used as a negative control. Mean and SEM are shown, data shown for n=5 from three donors (MDA-MB-468) and n=3 from one donor (A-431). Statistical analysis: unpaired t-test, adjusted p-values are shown. **(b)** Results of the ADCP assay on-chip showing the relative phagocytic index of macrophages (MФ) in MDA-MB-468 cells upon treatment with anti-EGFR IgA alone or in combination with the SIRPα-Fc, delivered via retargeted adenoviruses. Non-treated cancer cell-macrophage co-cultures were used as a control (MФ only); adenovirus encoding eGFP (mock) was used as a vehicle control. As a positive control for the maximum phagocytic activity, a combination of recombinant anti-EGFR IgA at 1 nM and SIRPα-Fc at 0.5 nM were used (prot. ctrl). Administration of the SIRPα-Fc alone was tested as an additional control condition. Mean, range, interquartile range and individual values are shown to indicate level of variability, data for at least n=5 from three different batches of monocyte-derived macrophages (from pooled donors). Statistical analysis: one-way ANOVA with Dunnett’s correction for multiple comparisons, adjusted p-values are shown. In both assays, MOI used: anti-EGFR IgA-encoding vector: 50 i.v.p./cell; SIRPα-Fc-encoding vector: 5 i.v.p./cell.

We tested the co-delivery of IgA and SIRPα-Fc in the ADCP on-chip assay in the MDA-MB-468 model (Figure 5b). Whereas the highest ADCP activity on-chip was observed for the control group to which recombinant proteins were added, significant activation of ADCP was only seen for AdV-mediated co-delivery of both payloads (on average a 2.3 ± 0.4-fold increase over the background), compared to 2.1 ± 0.5-fold increase for IgA alone, and 1.7 ± 0.7-fold increase for SIRPα-Fc alone.

### 2.6. IgA-based therapy leads to specific elimination of malignant cells in ADCC on-chip

An important aspect of any cancer therapy is its ability to selectively target cancer cells while leaving healthy cells unaffected. To evaluate the specificity of our adenovirus-mediated IgA therapy, we incorporated healthy dermal fibroblasts into our tumor-on-a-chip model, creating a tumor microenvironment containing both cancerous and non-cancerous cells. We then used this model to test the two most effective IgA treatments identified in our previous experiments: anti-EpCAM IgA in the HCT116 model and the combination of anti-EGFR IgA and the SIRPα-Fc protein in the MDA-MB-468 model. We assessed the ability of these treatments to activate neutrophils and kill the cancer cells while monitoring the impact on the healthy fibroblasts. In both models, we observed that the IgA-based treatments specifically targeted and eliminated the cancer cells while leaving the healthy fibroblasts unharmed (Figure 6a, b). This demonstrates the high specificity of our adenovirus-mediated IgA therapy, which can distinguish between cancerous and non-cancerous cells.

**Figure 6.**
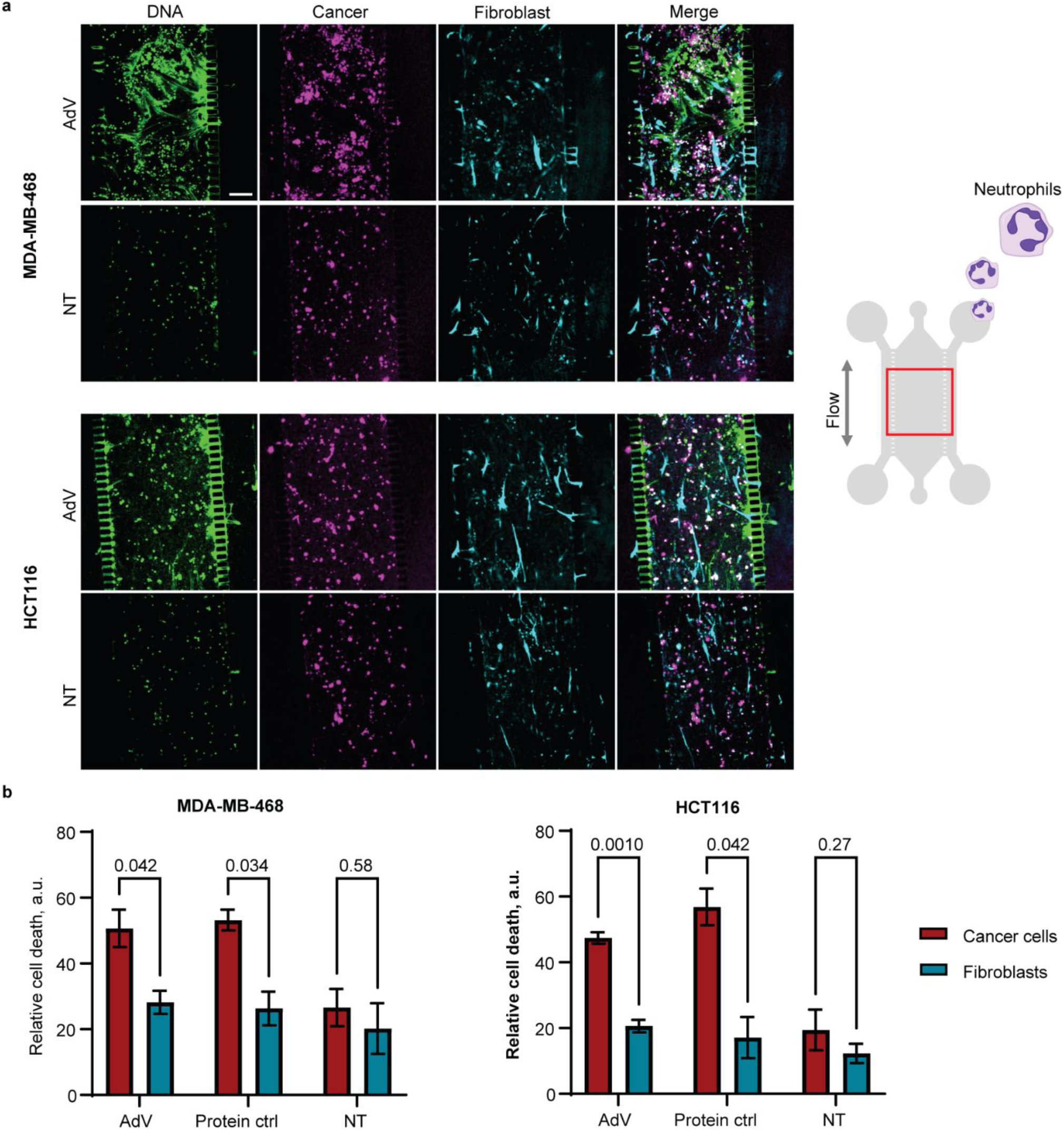
IgA-based, adenovirus-mediated treatment leads to specific elimination of cancer cells but not healthy fibroblasts in neutrophil ADCC on-chip. **(a)** Anti-EpCAM IgA, or anti-EGFR IgA together with SIRPα-Fc, were locally produced upon adenoviral transduction (AdV) in HCT116 and MDA-MB-468 tumor models, respectively. In each model, the respective cancer cells were labeled (magenta, CellTrace Far Red) and co-cultured on-chip with labeled healthy fibroblasts (cyan, CellTrace Violet). Following adenoviral transduction, the ADCC assay with primary human neutrophils was performed as above, and the cell death was detected using a DNA-binding dye (CellTox Green). The overlay images are shown demonstrating specificity for cancer cell killing compared to fibroblasts. The schematic image of the chip is shown with the approximate area imaged framed in red. As positive controls, recombinant proteins were perfused on-chip (Supplementary Figure S14). NT, negative control (non-treated co-culture with neutrophils added). Scale bar: 200 μm. Representative images are shown from four chips from two independent experiments (AdV samples: three chips). MOI: anti-EGFR IgA and anti-EpCAM IgA-encoding vectors: 50 i.v.p./cell; SIRPα-Fc-encoding vector: 5 i.v.p./cell. These MOI values were used in the functional assays as they achieved saturating signals of the payload expression. **(b)** Results of the ADCC assay on-chip, quantification performed per cell type for killing specificity. AdV, adenovirus-based delivery of anti-EpCAM IgA (in HCT116 model) or anti-EGFR IgA with SIRPα-Fc (in MDA-MB-468 model); Protein ctrl, respective recombinant protein controls perfused on-chip at 5 nM; NT, non-treated co-cultures to which neutrophils were added. Protein Ctrl (purified anti-EGFR IgA (left) or anti-EpCAM IgA (right)): n=2; AdV: n=3. Means with SEM are shown. Statistical analysis: unpaired t-test with the p-values adjusted for multiple comparisons indicated.

### 2.7. Effective production and therapeutic activity of IgA and SIRPα-Fc in tumor-bearing mice upon AdV injection

To validate the therapeutic potential of our adenovirus-mediated IgA therapy *in vivo*, we built upon the tumor-on-a-chip data and investigated the ability of the adenoviral vectors to induce local expression of IgA antibodies and the checkpoint blocker SIRPα-Fc in tumor-bearing mice. We used three different tumor models: MDA-MB-468 and HCT116 tumors, which were also used for tumor-on-a-chip experiments, were grown in immunocompromised SCID mice, and B16F10-EGFR tumors, which were used in immunocompetent C57BL/6 mice. B16F10-EGFR tumors grow slower in immunocompetent C57BL/6 mice as compared to parental B16F10 cells,^68^ but this *in vivo* model has nonetheless proven value for testing anti-EGFR therapies.^3^ This combined approach allowed us to evaluate the therapeutic approach in both the presence and absence of a functional immune system, and cross-validate results from our tumor-on-a-chip model (Figure 7a).

**Figure 7.**
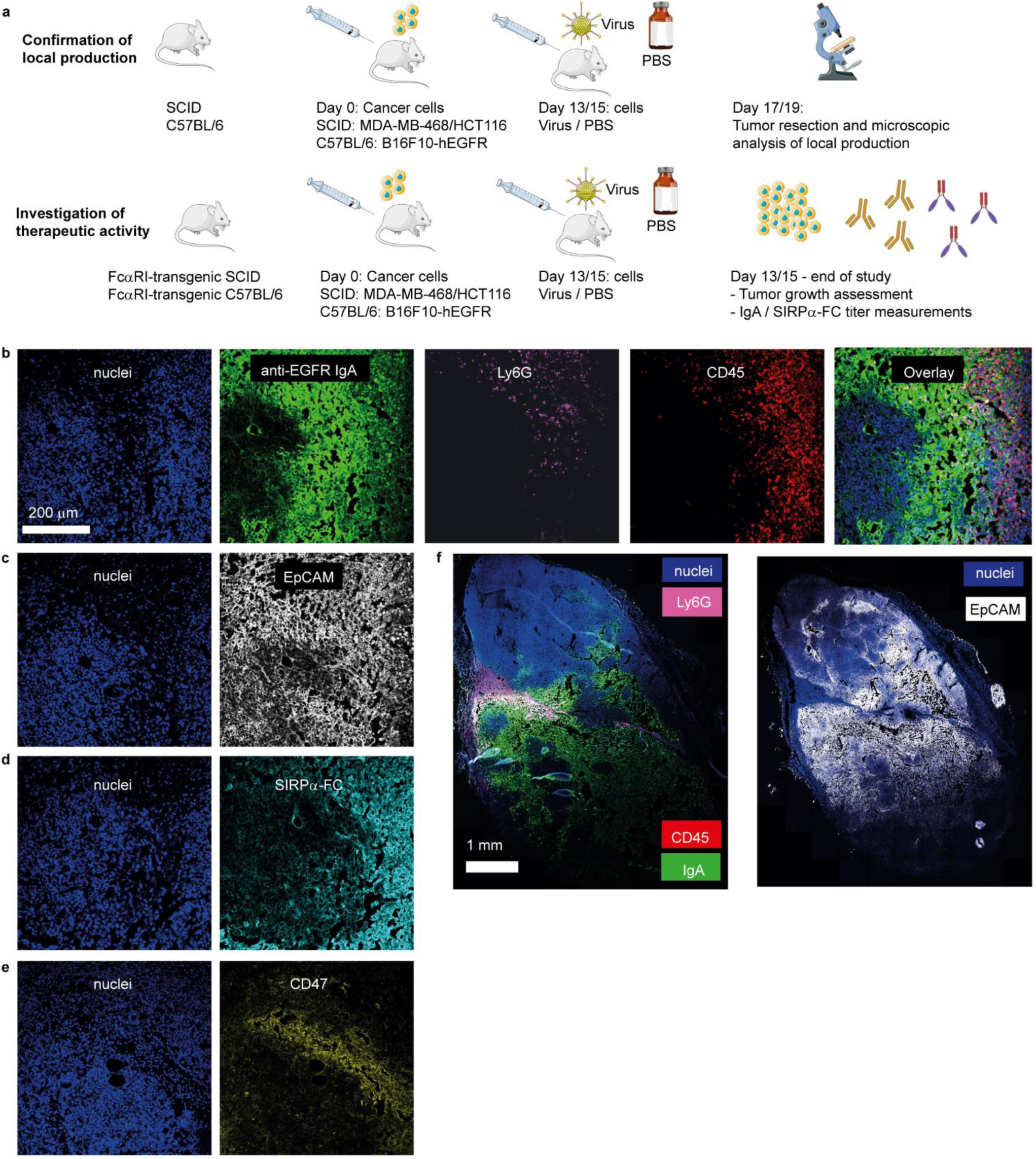
Immunofluorescence of MDA-MB-468 tumor slices from SCID mice injected with anti-EGFR IgA- and SIRPα-Fc-encoding adenoviral vectors. **(a)** Set-up of the in vivo study. **(b-e)** Immunofluorescence (IF) of tumor slices from MDA-MB-468 tumor-bearing SCID mice four days after injection with EpCAM-directed anti-EGFR IgA-encoding and SIRPα-Fc-encoding adenoviral vectors. Nuclei were stained with DAPI, and the produced anti-EGFR IgA was detected with a goat anti-Ckappa light chain. Ly6G and CD45 were detected in the same slice as markers for neutrophils and immune cells, respectively (**b**). EpCAM was detected to identify tumor cells with an anti-EpCAM antibody **(c)**, SIRPα-Fc with goat anti-human Fc gamma **(d)** and CD47 with an anti-CD47 antibody **(e)**. **(f)** A tilescan of a slice containing a full cross-section of the resected tumor was analyzed by IF for the condition described in (b) (left), or EpCAM (right), showing the distribution of the antibodies and infiltration of immune cells. The scale bar in (b) is applicable to all images from b-e. n = 6, representative images are shown.

We injected the adapter-carrying AdV vectors, which were additionally shielded from the immune system and thereby detargeted from the liver as reported before,^44^ directly into the tumors. We subsequently assessed the production and activity of the IgA antibodies and the SIRPα-Fc protein. We observed robust production and binding of anti-EGFR IgA (Figure 7b) to MDA-MB-468 tumors, which are both EpCAM- and EGFR-positive (Figure 7b,c) in SCID mice four days after a single injection. Similarly, SIRPα-Fc production and binding activity was robust in MDA-MB-468 tumors (Figure 7d) that were confirmed to be CD47 positive (Figure 7e). A high immune cell infiltration was also observed after a single injection (Figure 7b), as detected by increased staining of the pan-immune marker CD45. When looking specifically for neutrophils using the marker Ly6G, we also saw enrichment predominantly in areas with high payload presence. An analysis of a full cross-section of a slice of the resected tumor indicated that therapeutic antibody production occurred throughout most of the observable tumor, with neutrophil infiltration particularly strong at the edges of the tumor, indicating an active infiltration process is underway (Figure 7f).

No immune cell infiltration was seen in vehicle-treated mice (Figure S15). Similar findings were observed for EGFR-directed adenoviral vectors for the local production of anti-EpCAM IgA and SIRPα-Fc in HCT116 tumor-bearing SCID mice (Figure S16, Figure S17). For the immunocompetent B16F10 C57BL/6 model, the adenoviral viral vector and the produced IgA were both targeted to EGFR as B16F10 cells do not express human EpCAM. Also in this model, the local production of anti-EGFR IgA and SIRPα-Fc proteins was visible throughout most of the tumor (Figure S18), and a marked increase in immune cell infiltration upon treatment was seen (Figure S18, Figure S19).

To evaluate the therapeutic efficacy of our approach in vivo, we conducted further experiments in transgenic mice with a functional FcαRI receptor, which is necessary for IgA to activate immune cells.^41^ We compared the anti-tumor effects of AdV encoding different IgA formats (monomeric and dimeric) against distinct target antigens (EGFR and EpCAM), as well as the combination of monomeric anti-EGFR IgA with SIRPα-Fc.

In MDA-MB-468 tumor-bearing SCID mice, monomeric anti-EGFR IgA alone and in combination with SIRPα-Fc as well as dimeric IgA alone all significantly reduced tumor growth as compared to the vehicle control, with the anti-EGFR dimer and combination of monomer with SIRPα-Fc showing the strongest effects (55% and 57% reduction in tumor growth, respectively), 29 days after a single injection (Figure 8a). The retarded tumor growth led to a statistically significant improved survival only for the anti-EGFR dIgA condition, as evaluated using a Kaplan-Meier curve and a Log-rank test (Figure 8b). In HCT116 tumor-bearing mice, adenoviral vectors encoding the anti-EpCAM mIgA were compared with the anti-EpCAM dIgA and vehicle. In contrast to the MDA-MB-468 results, reduced tumor growth was only significant for the anti-EpCAM dIgA and did not reach significance for the anti-EpCAM mIgA, while no significant survival differences were observed for either condition (Figure 8c,d). In the immunocompetent B16F10 model in C57BL/6 mice, a strongly reduced tumor growth for the combination treatment was observed (Figure 8e), but this did not lead to a significantly increased survival (Figure 8f).

**Figure 8.**
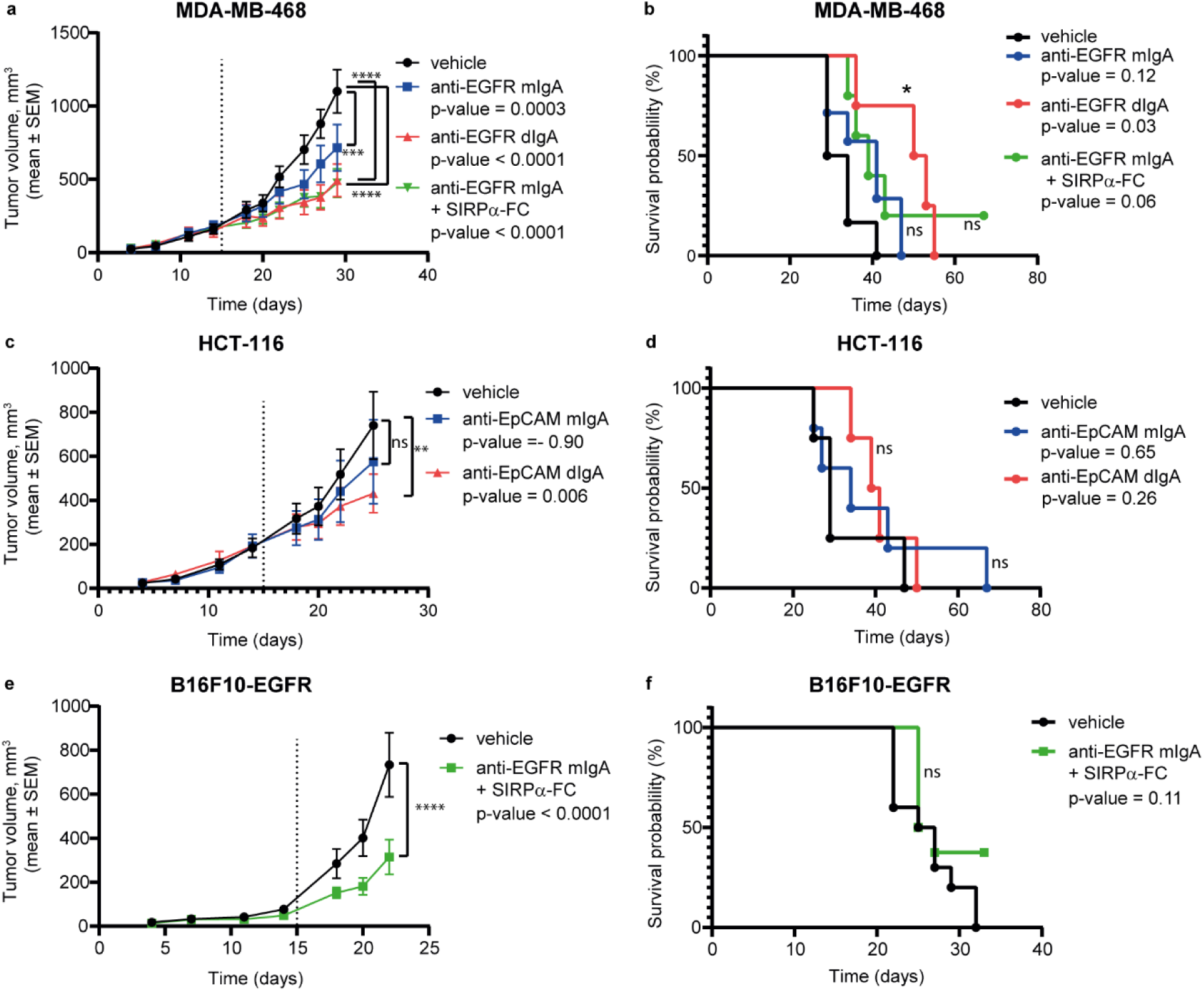
Tumor growth and survival curves of tumor-bearing FcαRI transgenic mice treated with adenoviral vectors encoding IgA and SIRPα-Fc proteins. **(a)** Tumor growth curves of MDA-MB-468 tumors in FcαRI-transgenic SCID mice treated with adenoviral vectors encoding the indicated proteins or vehicle control (PBS). Adenoviral vectors were retargeted to the MDA-MB-468 tumor cells using EpCAM-retargeted adenoviral vectors. **(b)** Kaplan-Meier curve illustrating survival of the treated mice as described in **(a). (c)** Tumor growth curves of HCT116 tumors in SCID mice treated with adenoviral vectors encoding the indicated proteins or vehicle control (PBS). Adenoviral vectors were retargeted to the HCT116 tumor cells using EGFR-retargeted adenoviral vectors. **(d)** Kaplan-Meier curve illustrating survival of the treated mice as described in **(c). (e)** Tumor growth curves of B16F10 tumors in immunocompetent FcαRI-transgenic C57BL/6 mice treated with adenoviral vectors encoding the indicated proteins or vehicle control (PBS). Adenoviral vectors were retargeted to the B16F10 tumor cells using EGFR-retargeted adenoviral vectors. **(f)** Kaplan-Meier curve illustrating survival of the treated mice as described in **(e).** Dotted lines indicate time of treatment. Statistical analyses of treatment-induced changes in tumor growth were done using a mixed model approach with a Geisser-Greenhouse correction. Asterisks indicate significant changes in tumor growth trajectories between treatment conditions and vehicle control (in **a,c,e**). Survival curves were analyzed using a Log-Rank test, with a Bonferroni correction applied for multiple testing. p values which were < 0.05 after correction for multiple testing were considered significant. Testing was done vs. vehicle for all conditions.

We also measured the levels of IgA and SIRPα-Fc in the blood of treated mice. Monomeric IgA yielded measurable titers in most FcαRI-transgenic mice, with 6 out of 7 in the MDA-MB-468 model (Figure S20) and 5 out of 7 in the HCT-116 model (Figure S21) in conditions where only the monomeric IgA-encoding adenovirus was administered. In the immunocompetent B16F10 FcαRI-transgenic model, where only the combination of anti-EGFR IgA and SIRPα-Fc was tested, IgA titers were generally lower and detectable in 4 out of 12 mice (Figure S22), in line with the trend for the combination in the MDA-MB-468 model (Figure S20). Dimeric IgA titers were either undetectable or very low (Figure S20 and S21), presumably in part due to a lower degree of intravasation into the bloodstream because of their larger size, and in part because they are cleared more rapidly from the circulation due to an enhanced recognition by phagocytic cells, particularly in the liver and spleen. Alternatively, dimeric IgA may bind to pIgR and be transported across epithelial cells to mucosal surfaces, such as the respiratory tract, gastrointestinal tract and genitourinary tract^69^. SIRPα-Fc remained below detection levels at all time points, possibly due to binding to the abundant amounts of endogenous murine CD47 (data not shown). Finally, we analyzed the expression of immune-related genes in the tumors of surviving C57BL/6 FcαRI transgenic mice after 32 days. However, we did not observe any significant differences between treated and untreated mice at the time point analyzed (Figure S23).

## 3. Discussion

In this study, we demonstrate the feasibility of a novel approach to cancer immunotherapy: using retargeted adenoviruses to convert tumor cells into "biofactories" for the localized production of IgA antibodies and an innate immune checkpoint blocker. This strategy focuses on engaging myeloid cells, key players in the innate immune response, to effectively target and eliminate cancer cells. Our findings highlight the potential of this approach to overcome the challenges associated with systemic delivery of IgA and to enhance anti-tumor immune responses, and identifies dimeric IgA both *in vitro* and *in vivo* as the preferred candidate. Of note, use of dimeric IgA necessitates a local production approach as it would be very rapidly removed from the circulation when injected intravenously.

We engineered a multicistronic vector system for efficient production of IgA antibodies, achieving comparable functionality to previously reported systems.^5,51,70^ Interestingly, we observed a slightly higher molecular weight for the IgA heavy chains produced from our system, which might be attributed to differences in glycosylation patterns due to different production systems.^71^ Importantly, our tricistronic vector system allowed for efficient production of dimeric IgA, achieving up to 50% dimer fraction without the complex and time-consuming purification steps that are required for pure dimeric IgA production.^22,72^

We confirmed the ability of our system to produce and distribute IgA antibodies and the SIRPα-Fc checkpoint blocker within the tumor microenvironment, both in a tumor-on-a-chip system and *in vivo*. The ability to locally secrete therapeutic proteins upon viral delivery is crucial to reach most or all target cells, and thus for maximizing therapeutic efficacy, as adenoviral transduction efficiency is often limited in 3D environments and *in vivo.*^47,73–75^

Our findings align with previous research demonstrating the potential of adenovirus-mediated gene therapy to induce transduced cells to locally secrete therapeutic agents. This approach is particularly promising for delivering synergistic combinations of multiple therapeutics simultaneously, as recently shown by Brücher *et al.*, who used retargeted adenoviral vectors to co-deliver an anti-PD1 antibody, interleukin (IL)-12, and IL-2 to cancer cells.^45^ To avoid potential competition between the adenovirus and the produced IgA for the same receptor, and to facilitate repeated administrations,^73^ we targeted the adenovirus to a different receptor than the IgA. For example, we delivered anti-EGFR IgA using an EpCAM-targeted adenovirus. However, in the B16F10-EGFR mouse model, which lacks human EpCAM, both the adenovirus and the IgA were targeted to EGFR. Since our focus was on the delivery approach and antibody framework, we selected the well-established targets EGFR and EpCAM, which nonetheless also have some expression on healthy epithelium that could give rise to some on target toxicities, which our study cannot rule out.

To gain a deeper understanding of IgA-mediated tumor cell killing through our approach, we used microfluidic tumor-on-a-chip models. These models provide a more physiologically relevant environment than traditional 2D cell cultures, allowing for detailed analyses of immune cell behavior within a 3D tumor microenvironment, including cell growth in the ECM, interactions with an ECM-medium interface, response to chemoattractant gradients, and immune cell infiltration and migration. Using these models, we observed potent activation of myeloid immune cells by anti-EpCAM and anti-EGFR IgA, leading to ADCC and ADCP of cancer cells, in line with previous reports for anti-EGFR IgA activity *in vitro.*^5,25^ We demonstrate significant activation of anti-tumor neutrophil ADCC activity, particularly for dimeric IgA, as seen before,^22^ and this was achieved with IgA produced by the transduced tumor cells. Importantly, results from these models correctly predicted the subsequent observations *in vivo* with respect to treatment-induced neutrophil and CD45+ immune cell infiltration and therapeutic efficacy. The local production approach overcomes bottlenecks associated not only with production, but also with systemic delivery, such as a short plasma half-life^76^ and the need to extravasate to reach the tumor. Noteworthy, our dimeric IgA was engineered based on the modified IgA2.0 framework^5^ with re-introduced Cys-Tyr residues, allowing binding of the J-chain, but it may have a reduced ability to stably interact with the secretory component (SC), which underlies IgA dimer transcytosis, due to the Cys to Ser mutation at position 311.^77,78^ Intriguingly, Biswas *et al*.^79^ found that co-localization of (secretory) IgA but not IgG with pIgR in high-grade serous ovarian cancer samples was associated with a favorable clinical outcome. Their investigation *in vitro* suggested the possible mechanism to be secretory IgA-mediated downregulation of tumor-promoting ephrins, upregulation of inflammatory genes and inhibitors of the RAS pathway, and compromised MAPK signaling,^79^ thus highlighting a promising future research direction that harnesses the ability of dimeric IgA to bind the secretory component while leveraging our localized production approach. Interestingly, our tumor-on-a-chip model also revealed differences in the concentration necessary to shown an effect of IgA compared to traditional 2D models, with higher concentrations required in the tumor-on-a-chip model, highlighting the importance of using more physiologically relevant systems for preclinical testing.^33,80^ We also observed the formation of neutrophil extracellular traps (NETs), web-like structures of filamentous DNA that also contain various proteins, in response to IgA-mediated activation, likely due to potent FcαRI activation.^66,67^ While NETs have traditionally been associated with pro-tumorigenic effects,^81–83^ our findings suggest a context-dependent role for NETs in tumor progression. The observation of filamentous DNA structures, which we presume to be NETs, occurring specifically in areas with high IgA-mediated tumor cell killing, suggests that activated neutrophils can utilize NETosis as an anti-tumor mechanism,^82,84^ though it could also be a co-occurring event not directly related to therapeutic efficacy.

In addition to ADCC, we showed ADCP by macrophages to be another effector mechanism of cancer cell elimination in response to adenovirus-based IgA delivery. As shown before with recombinant IgA, M2-macrophages can be activated to eliminate cancer cells,^33^ which was also demonstrated *in vivo.*^21,33^ In our on-chip ADCP assay, *in situ* produced anti-EGFR and anti-EpCAM IgA induced significant phagocytic activity by M2-activated macrophages. The effects were restricted to a severalfold increase in ADCP activity, particularly because we already noticed substantial anti-tumor effects of non-stimulated M2 macrophages in our microfluidic system. Nonetheless, Our results further strengthen the therapeutic potential of our approach, as it engages multiple effector mechanisms of the myeloid immune system. One limitation of our current study is an inaccurate representation of the immunosuppressive tumor-associated macrophage phenotype that is typically observed *in vivo* which, when incorporated, is expected to provide a better pathophysiological relevance of the tumor model.

We furthermore show how our approach can be extended for co-delivery of an innate checkpoint blocker to boost effects of IgA-mediated ADCC and ADCP and specifically eliminate cancer cells. To block CD47, we chose SIRPα-Fc with the Fc modified to ablate effector functions, as this approach was reported earlier to boost IgG^34^ or IgA^28^ effects *in vivo*. We demonstrate how our local production system of both IgA and SIRPα-Fc can be used to achieve the reported effect using an approach that can in principle overcome the need for repeated administrations^2,76^ and possible systemic toxicity risks. In our study, even after a single adenovirus injection, we could see long-term benefit with respect to tumor growth, which is consistent with the long-term production of IgG antibodies until day 61 as seen after intratumoral injection with an IgG-producing adenovirus.^46^

Our study contains several limitations. For capacity reasons and to limit animal usage, we did not include IgG benchmarks in our in vivo studies, meaning a direct comparison of activity against similar IgG isotype-based molecules cannot be done. However, others have extensively performed such studies in recent years for a variety of targets including EGFR, HER2, CD20 and GD2.^3,27,29,30^ When correcting for pharmacokinetic effects, anti-tumor effects of similar magnitude as for IgG counterparts were generally found, yet through distinct immunological pathways. For GD2, next to comparable tumor clearance a striking reduction in neuropathic pain was observed with IgA, due to reduced complement activation and inflammation.^30^ By not including IgG isotype antibodies, we could also not study potential synergistic effects, which we hypothesize may exist due to activation of different pools of immune effector cells, as shown by others, for instance through hybrid IgA/IgG Fc tails or combination therapeutic approaches.^85–87^ In our *in vivo* studies, we did not make comparisons with vectors encoding irrelevant proteins (e.g. GFP) as controls, meaning the observed effects compared to PBS injection controls are due to the local expression of therapeutic molecules and the biological effects of the vector itself. We did include such control vectors in our organ-on-a-chip system. In the *in vivo* studies, our focus and hence experimental design was optimized for comparing activity of the expressed proteins (e.g. monomeric vs. dimeric vs. combination), for which the data are robust. The *in vivo* effects of the viral vector itself have been characterized in previous studies by us.^44,46,88^ Mechanistically, we focused mostly confirming successful therapeutic protein production in non-transgenic mice. Our studies in transgenic mice were set up to detect effects on tumor growth, which is why expression of immune-related genes was only studied at day 32, at which point no changes in expression levels were seen. In future studies, it would be interesting to also look at cytokine levels and immune cell infiltration at early timepoints in transgenic mice models.

We also did not study the adaptive arm of the immune system. Yet, as others have provided evidence of the ability of neutrophil-mediated tumor cell killing^85^ and blockade of the CD47-SIRPα axis^34^ to enhance the adaptive arm of the anti-tumor response, it will be interesting to investigate in more depth how our approach can stimulate anti-tumor adaptive immune responses in longer term *in vivo* immunocompetent studies, possibly aided by a localized production of an immunostimulatory agent and/or the use of a hybrid IgA/IgG Fc framework. Furthermore, while we observed promising anti-tumor effects *in vivo* and could robustly detect locally produced monomeric IgA, the levels of dimeric IgA and SIRPα-Fc in the bloodstream were low or undetectable. This could be due to several factors, including more rapid clearance of the dimeric IgA due to an enhanced recognition by phagocytic cells, particularly in the liver and spleen, transport to mucosal surface via pIgR binding, better target binding, and the localized nature of our delivery approach where produced proteins may be degraded before reaching the circulation, and is in line with a reduced risk of systemic side effects.

Recently, adenoviral vectors have received increasing attention due to their promise of utilization in cancer immunotherapy, particular given their high capacity that enables the delivery of multiple genes simultaneously for effectively reprogramming cold tumor microenvironments. However, challenges still need to be overcome, in particular the pre-existing immunity to adenoviruses and potential inflammatory responses in non-target tissues such as the liver, spleen and kidney, which increase translational challenges. Further advancements in vector engineering are underway to overcome these obstacles, paving the way towards clinical applications.^89,90^

Altogether, we show for the first time how the therapeutic potential of IgA molecules as cancer-fighting agents can be realized while overcoming challenges of its systemic delivery using the retargeted adenovirus platform. Our strategy thus highlights an alternative approach towards cancer immunotherapy based on IgA and a local production, which may potentially work synergistically with IgG-based therapies by engaging different effector mechanisms, as also exemplified by recent work using IgG/A cross-hybrid molecules.^85–87^ Next steps in the development of our approach involve expansion to a broader range of biologics for co-delivery via retargeted adenoviruses (e.g. cytokines, IgG), and development towards high-capacity, “gutless” adenoviral vectors,^45,47,91^ which is hypothesized to lead to more complete and more sustained anti-tumor responses. This will then pave the way for novel therapeutic modalities to convert tumors into therapeutic “biofactories” for long-term potent anti-cancer benefits.

## 4. Methods

### 4.1. Cell culture

MDA-MB-468 (breast carcinoma), HCT116 (colorectal carcinoma), A-431 (epidermoid carcinoma), MCF-7 (breast carcinoma), OVCAR3 (ovarian carcinoma), MC38 (C57BL6 murine colon adenocarcinoma) and HEK293 (human embryonic kidney) cell lines were obtained from ATCC. Primary human dermal C5120 fibroblasts were a kind gift of Dr. Rodenburg (Radboud University Medical Center, the Netherlands) and have been extensively characterized before^92^. The generation of B16F10-luc-EGFR cells (murine melanoma) was reported before.^3^ FreeStyle™ Chinese Hamster Ovary Cells (CHO-S, Thermo Fisher Scientific, Dreieich, Germany) were used for production of recombinant proteins. Human umbilical cord endothelial cells (HUVECs) were obtained from Lonza (Basel, Switzerland) and used until passage 10. The cancer cell lines and HEK293 cells were maintained in Dulbecco’s Modified Eagle Medium (DMEM, Gibco, Amarillo, TX, USA) supplemented with 10% fetal calf serum (FCS, PAN-Biotech, Aidenbach, Germany). C5120 fibroblasts were maintained in Medium 199 (Gibco) supplemented with 10% FCS. HUVECs were maintained in Endothelial Cell Growth Media-2 (basal medium EBM-2 (Lonza) supplemented with the BulletKit supplement (Lonza). FreeStyle CHO cells were maintained in suspension in CHOgro expression medium (Mirus Bio, Madison, Wisconsin, USA) supplemented with 20 mM glutamine and 0.3% poloxamer 188 (Mirus Bio), on an orbital shaker at 180 rpm (diameter 50 mm). For on-chip experiments, media for tumor cells and C5120 were further supplemented with 1× penicillin-streptomycin (Pen-Strep, Merck, Darmstadt, Germany) and amphotericin B (1 μg/mL, Merck). Adherent cells were cultured at 37°C in a humidified incubator with 5% CO_2_, referred to as standard cell culture conditions. CHO-S cells were cultured in a humidified incubator with 8% CO^2^ at 37°C for maintenance, and at 31°C for production of recombinant proteins.

### 4.2. Assembly of the pShuttle expression system

The pShuttle vector engineered for IgA and SIRPα-Fc production was adapted from the AdEasy Adenoviral Vector System (Agilent Technologies, Paris, France) and the previously reported SHREAD system.^46^ The expression cassettes for IgA2.0^5^ and SIRPα-Fc^34^ were synthesized by Twist Bioscience (San Francisco, CA, USA) and cloned into the pShuttle vector via Gibson assembly (New England Biolabs, Ipswich, MA, USA). The open reading frame (ORF) under a CMV promoter consisted of a rituximab VH leader sequence,^93^ VH (heavy chain variable domain)-CH1-hinge-CH2-CH3 (heavy chain constant domains) heavy chain cassette, followed by a furin recognition site-GSG-P2A or T2A self-cleaving peptide sequence for efficient separation of the IgA chains, followed by a rituximab VL leader sequence^93^ and VL (light chain variable domain)-Ckappa (light chain constant domain) with a BGH poly-A signal downstream. The expression system included restriction sites for easy exchange of expression cassettes (Figure S1). The anti-EGFR IgA2.0 heavy and light chain single ORF expression cassette was cloned following PCR amplification with generic primers containing a 5’-GTGAACCGTCAGATCCGC-3’ upstream overhang and 5’-GATCCGGTGGATCGGATATC-3’ downstream overhang for the Gibson assembly. The construct was cloned into the pShuttle backbone via *NheI-HindIII* restriction sites.

For multicistronic expression of the IgA heavy and light chains within the single ORF, ribosome skipping-inducing elements P2A (derived from porcine teschovirus-1 2A) or T2A (derived from *thosea asigna* virus 2A) were used, as reported earlier.^70^ The P2A or T2A sequences contained an N-terminal GSG-spacer to improve the sequence translation efficiency. To enable *in situ* removal of the 2A peptide sequence from the C-terminus of the heavy chain after expression, a furin cleavage site was used upstream of the GSG-2A sequence.^45^ The P2A or T2A sequences were exchanged in the pShuttle vector via *SalI-KpnI* restriction enzymes and Gibson assembly with the overlap sequences 5’-CATGTCAATGTGTCTGTTGTCATG-3’ and 5’-GTTATTCAGCAGGCACACAACAG-3’ upstream and downstream, respectively.

The anti-EpCAM IgA2.0 was engineered by grafting the VH and the VL domains from the previously reported humanized anti-EpCAM single-chain Fv fragment (scFv)^52^ on the IgA2.0 framework via Gibson assembly: The heavy chain variable fragment was cloned via *NheI-XhoI* restriction sites with the 5’-GTGAACCGTCAGATCCGCTAGAGATCTACAGCTAGCGCCGCCACCATG-3’ upstream and 5’-TTCTGAACCTAAGAGCAGGTCCTCGAGG-3’ downstream overlaps; the light chain variable fragment was subsequently cloned via *SalI-KpnI* restriction sites as described above.

To generate dimeric IgA2.0 expression constructs, the residues Cys-Tyr were reintroduced at the C-terminus into the heavy chain fragment to enable disulfide bond formation between the joining chain (J-chain) and heavy chain. The EMCV IRES was placed in front of the J-chain leader sequence followed by a His6 tag and the J-chain, and this unit was introduced downstream of the light chain sequence to express J-chain with the multicistronic heavy chain-light chain cassette. The construct was cloned via the *AgeI* restriction site following PCR amplification with 5’-GAGCTTCAACAGGGGAGAGTGTTGATGACC-3’ and 5’-ACAACAGATGGCTGGCAACTAGAAGGCACAG-3’ and Gibson assembly.

The SIRPα-Fc CD47 blocker sequence was composed of the SIRPα leader sequence, the consensus domain 1 with an N80A mutation of the D1 domain of SIRPα reported earlier^34^ and the Fc part with the mutations N297A, S298A, E333A, and K334A (termed AAA-N297A mutant)^54^. The fragment containing the EF1a promoter with the expression cassette was cloned into the pShuttle vector by Gibson assembly with the upstream 5’-TTACTCATAGCGCGTAATACTGGTAC-3’ overlap and downstream 5’-CCTTAACCACGCCCAGATCAAGCTTCTACGATACCGATAGAGATGGG-3’ overlap. The expression vectors were sequence-validated by Sanger sequencing of the inserts.

### 4.3. Adenoviral vector production and purification

The adenovirus generation was performed according to the AdEasy Adenoviral Vector System manual (Agilent Technologies) using the modified pAdEasy-1-HVR7 adenoviral genome^46^ and using HEK293 cells as a packaging cell line. The viral particles were purified using a two-step cesium chloride density gradient ultracentrifugation, and dialyzed against 20 mM HEPES pH 8.1, 150 mM NaCl, 1 mM MgCl_2_. Glycerol was added to a final concentration of 10% (v/v) and viral vectors were snap-frozen, and stored at -80°C. The virus titers were determined as optical viral particles (o.v.p.) or infectious viral particle (i.v.p.), used later for the calculation of the respective multiplicity of infection (termed MOI or infectious MOI, respectively) per cell. For the o.v.p. determination, the absorbance was measured at 260 nm using a Nanodrop One UV Spectrophotometer (Thermo Fisher Scientific) as reported earlier.^46^ For the i.v.p. titer determination, quantitative PCR was performed as reported earlier.^46^ For the retargeting of adenoviral particles to EGFR or EpCAM, trimeric knob adapters were produced and purified in *E. coli* as previously described.^43^

### 4.4. Production and purification of recombinant IgA and SIRPα-Fc proteins

Recombinant IgA and SIRPα-Fc proteins were produced using the pShuttle vector in suspension FreeStyle™ CHO-S cells as follows: the cells were seeded at a density of 3×10^6^ cells/mL 24 h prior to transfection. On the day of the transfection, the cell density was adjusted to 4×10^6^ cells/mL in fresh medium, and the transfection mix (total of 2.5 mg plasmid DNA per 1 L culture, complexed with linear polyethyleneimine-20,000 (Merck) at a 1:2 ratio) was added. The cell suspension was supplemented with 0.5 mM valproic acid (Mirus Bio) 24 h post transfection. The cells were incubated at 31°C, 8% CO_2_ while shaking, and the supernatants were harvested after 7 days. For the purification of IgA, the supernatants were sterile-filtered through a 0.2 μm filter, loaded onto a 1 mL HiTrap Protein L prepacked column (Cytiva, Grens, Switzerland), washed with 10 column volumes of PBS, and the antibodies were eluted with 5 column volumes of 0.1 M glycine, pH 3.0, and neutralized with 375 µL of 1 M Tris, pH 8.8. The elution fractions with detectable IgA were pooled and concentrated using Amicon Ultra centrifugal filter units (Merck) with a molecular weight cutoff (MWCO) of 50 kDa. The samples were desalted, and the buffer was exchanged into endotoxin-free Dulbecco’s PBS (DPBS, Millipore Merck). The concentration of IgA was measured at 280 nm using a Nanodrop One UV Spectrophotometer. The IgA proteins were stored at 4°C. The SIRPα-Fc protein was purified using Protein A MagBeads (Genscript, Piscataway, NJ, USA): the beads were loaded overnight at 4°C, washed with 0.1% (v/v) Tween-20 in PBS (PBS-T), and the protein was eluted with 0.5 mL of 0.1 M glycine, pH 3.0, and neutralized with 80 µL of 1 M Tris, pH 8.8. The elution fractions were pooled, and the protein was concentrated using Amicon ultracentrifugal filter units with a MWCO of 30 kDa. The samples were desalted, and the buffer was exchanged into the endotoxin-free DPBS. The protein concentration was measured at 280 nm using Nanodrop One UV Spectrophotometer, and the protein was stored at 4°C. Purified proteins were further analyzed for monomeric or dimeric structure with size exclusion chromatography using GE Superdex 200 increase 10/300 GL column (Cytiva).

### 4.5. Western blots

For the analysis of proteins under reducing conditions, protein samples were incubated in the presence of Laemmli buffer (Bio-Rad, Hercules, CA, USA) with β-mercaptoethanol as a reducing agent, at 96°C for 10 min. For the analysis under non-reducing conditions, the protein samples were combined with Laemmli buffer without β-mercaptoethanol at room temperature. The samples were loaded on Bolt 4-12% Bis-Tris gels (Thermo Fisher Scientific) in a 1× 3-(N-morpholino) propanesulfonic acid (MOPS) buffer. Chameleon Duo Pre-stained Protein Ladder (Li-Cor, Lincoln, NE, USA) was used as a molecular weight reference. The samples were separated at 150 V for 10 min followed by 200 V for 20 min. The proteins were then transferred on the TransBlot Turbo PVDF-LF transfer membrane (Bio-Rad) using a TransBlot transfer system according to the manufacturer’s instructions. The blocking was performed by incubating the membrane in a 50 mL centrifuge tube with 1× casein solution (Sigma, Steinheim, Germany) in PBS on a roller shaker for 30 min at room temperature in the dark. After blocking, the membrane was incubated with a primary antibody solution in 1:1 PBS-T/blocking solution, overnight at 4°C in the dark. The following antibodies were used: for IgA light chain detection: goat anti-human kappa chain (#2060-01, Southern Biotech, Birmingham, AL, USA, 1:2,500), or mouse anti-human kappa chain (#K4377, Sigma, 1:2,500 dilution); for IgA heavy chain detection: goat anti-human α heavy chain (#5104-2004, Bio-Rad, 1:2,500); for J-chain detection: mouse anti-poly His-tag (#H1029, Sigma, 1:2,500 dilution); for SIRPα-Fc detection: mouse anti-γ heavy chain (#2040-01, Southern Biotech, 1:2,500 dilution). The membranes were washed 3× 10 min with PBS-T and incubated with a secondary antibody solution (in 1:1 PBS-T/blocking solution) at room temperature for 40 min. The following secondary antibodies were used: donkey-anti-mouse IRDye800 (#926-32212, Li-Cor), donkey-anti-goat IRDye680 (#926-68074, Li-Cor) alone or in combination for multiplex detection. Membranes were washed 3 × 10 min with PBS-T and 1 × 10 min with PBS, and imaged using an Odyssey DLx Fluorescence Imaging System (Li-Cor).

### 4.6. ELISA

Nunc MaxiSorp flat-bottom 384 well plates (Thermo Fisher Scientific) were coated with 25 μl/well of 100 nM NeutrAvidin (Thermo Fisher Scientific) solution prepared in a carbonate buffer (pH 9.5), and incubated overnight at 4°C. All incubation steps were performed on an orbital shaker (2.5 mm diameter) at 700 rpm. The next day, the coating solution was aspirated and the plate was washed 3 × 5 min with PBS. The plate was coated with 25 μl/well of 50 nM EGFR ectodomain protein produced and purified as described,^94^ biotinylated EpCAM ectodomain protein (Acro Biosystems, Newark, DE, USA) or 0.25 mg/mL biotinylated CD47 protein (Sino Biological, Beijing, China) diluted in PBS, and incubated for 1 h at room temperature. The coating solution was aspirated, and the wells were blocked with 25 μl/well of 5% (w/v) BSA in PBS for 1 h at room temperature, followed by 3 × 10 min washing with PBS-T. The titrated standards (cetuximab, or anti-CD47 IgG control antibody (#11-0479-42, eBioscience, San Diego, CA, USA), or purified anti-EGFR IgA2.0, anti-EpCAM IgA2.0, or SIRPα-Fc), blanks, or the sample dilution series (cell culture supernatants diluted in 0.1% (v/v) Tween-20 + 0.5% (w/v) BSA in PBS (PBS-TB) were added in duplicate, 25 μl/well. The samples were incubated for 1 h at room temperature, followed by a 3 × 10 min washing with PBS-T. For the IgA or SIRPα-Fc detection, primary antibody (goat-anti-human kappa light chain #2060-01, Southern Biotech, or goat-anti-human IgG #2048-01, Southern Biotech) was diluted 1:2,000 as above, 25 μl/well was added and left to incubate for 1 h at room temperature. The plate was washed as above. The secondary antibody (alkaline phosphatase (AP)-conjugated rabbit anti-goat IgG (#A16145, Thermo Fisher Scientific) was diluted 1:2,000 in PBS-TB, added at 25 μl/well and incubated for 1 h at room temperature. The plate was washed as above. For the signal detection, 1 mg/mL solution of para-nitrophenyl-phosphate (4-NPP) substrate (Thermo Fisher Scientific) prepared in 4-NPP buffer (50 mM NaHCO_3_, 50 mM MgCl_2_), was added at 25 μl/well and incubated for 15 min at room temperature. The signal was measured with a VICTOR Nivo Multimode Microplate Reader (PerkinElmer, Waltham, MA, USA) by recording absorbance at 405 nm. Once the optimal signal was reached, the reaction was stopped with 3 M NaOH. The absorbance normalized to the reference wavelength was further normalized to the blank wells. The data was analyzed using GraphPad Prism Software v.9.5.1 for Windows (GraphPad Software, San Diego, CA, USA).

### 4.7. Microfluidic chip fabrication and loading

Master molds and replicas of the microfluidic devices were produced using SU8 photoresist and polydimethylsiloxane (PDMS) as described before^92^. The inlet and outlet holes for each of the three channels were made at a 45° angle using a 1.5 mm biopsy puncher (Kai Industries, Tokyo, Japan), and the medium reservoirs were made with a 5 mm biopsy puncher partially overlapping with the respective 1.5 mm holes to facilitate medium loading. The PDMS replicas were then bonded onto a high precision cover glass no. 1.5 (#0107242, Superior Marienfeld, Lauda-Königshofen, Germany) by oxygen plasma treatment in a plasma cleaner (PDC-32G-2, Harrick Plasma, Ithaca, NY, USA) for 1 min, and subsequent incubation on a hotplate for 20 min at 80°C. The devices were kept at least overnight prior to coating to avoid defects in bonding. Prior to cell loading, the chips were freshly coated with a polydopamine solution to enhance collagen adhesion (2 mg/mL dopamine hydrochloride (Sigma) in 10 mM Tris-HCl, pH 8.5) for 1 h at room temperature in the dark. The coating solution was removed, and the chips were thoroughly washed with milliQ followed by washing with 70% v/v ethanol under aseptic conditions. The excess ethanol was aspirated, the chips were placed in a Petri dish, dried for three hours at 65°C and kept sterile until loading. Cells were loaded into the tissue (middle) compartment in collagen (#354249 Rat tail collagen type I, Corning, New York, NY, USA) prepared as a 4 mg/mL working solution by neutralization with 37 g/l NaHCO_3_ and 1 M HEPES buffer combined in a 1:1 ratio, yielding a pH value of ∼7.4. The collagen solution was mixed with cells in the appropriate cell culture medium to the final 3 mg/mL collagen concentration and a cell density of 1 × 10^4^ – 6 × 10^4^ cells/chip (1.7 × 10^3^ – 1 × 10^4^ cells/µL). The collagen was gelated by incubation of chips at 37°C for 20 min. The cell culture medium was added into each of the side channels, 50 µL per each reservoir, and the chip was perfused by incubation under standard cell culture conditions on a Perfusion Rocker Mini (Mimetas, Oegstgeest, The Netherlands) to provide a bidirectional perfusion flow, at a 7° incline, 10 min per cycle. Unless stated otherwise, the on-chip culture medium was refreshed every 3 days.

### 4.8. Adenovirus-mediated gene delivery in monolayer and on-chip (co-) cultures

One day prior to viral transduction, the cancer cells were optionally labeled with CellTrace (Thermo Fisher Scientific) cytoplasmic dyes according to the manufacturer’s protocol, and seeded in an 18-well chamberslide (#80826, ibidi, Gräfelfing, Germany) at a density of 1 × 10^4^ cells/well. After an overnight incubation, transduction with retargeted AdV5 vectors encoding IgA and/or SIRPα-Fc was performed. On the day of viral transduction, virus-adapter complexes were prepared as described^92^. The AdV-adapter complexes in fresh medium were added to the cells, and incubated for 3 h. Following the incubation step, the medium with the non-adhered / non-internalized AdV complexes was removed, the cells were carefully washed three times with the cell culture medium. The cells were incubated for two more days to reach high expression levels of the AdV payloads (IgA and SIRPα-Fc). The AdV-mediated production of IgA or SIRPα-Fc proteins was confirmed with immunofluorescence (IF) (see below). For the on-chip gene delivery, cells were loaded on-chip and incubated overnight on the perfusion rocker. AdV-adapter complexes were formed as described and diluted in a total of 200 µL cell culture medium per each on-chip culture. The old medium was removed and the medium containing AdV-adapter complexes was added at 50 µL per each of the four medium reservoirs. The chips were incubated on a perfusion rocker, and after an overnight incubation the medium containing the non-internalized particles was replaced with fresh medium. The chips were then incubated for two more days. The AdV transduction on-chip was followed by IF on-chip to confirm the payload delivery and production, or in immunological assays (see below).

### 4.9. Immunofluorescence in monolayer and on-chip (co-)cultures

Immunofluorescence-based detection of IgA and SIRPα-Fc proteins in monolayer cultures was performed in ibidi μ-Slide 18-well chamber slides. Cells were optionally pre-labeled with different CellTrace cytoplasmic dyes, which enter cells as nonfluorescent esters, are converted to a fluorescent derivative by cellular esterases and subsequently react with protein amino groups via the N-hydroxy succinimidyl ester group. Cells were seeded at a density of 1-5 × 10^5^ cells/well. Following a 24-h incubation, the cells were washed with PBS. For the detection of IgA or SIRPα-Fc proteins in cell culture supernatants upon transfection with pShuttle expression vectors, the supernatants were diluted in PBS and incubated with the cells for 1 h. For detection of *in situ* produced IgA or SIRPα-Fc, first the AdV transduction was performed as described above, and then the cells were incubated for three days. On the day of the assay, the cells were washed with PBS, and the pure proteins were added as positive controls. As a positive control for SIRPα-Fc in the supernatants of the transfected cells, mouse anti-human CD47 antibody (#11-0479-42, eBioscience) was used at 0.25 μg/mL, and as a control for anti-EGFR IgA detection cetuximab (Merck) was used at 2.5 µg/mL in the cell culture medium. As a positive control for *in situ* produced IgA or SIRPα-Fc, the respective in-house made and purified respective proteins were used at 50 nM. The cells were incubated with the proteins for 1 h. The samples were washed with PBS and fixed with 4% (v/v) paraformaldehyde (PFA) in PBS for 15 min. The samples were blocked with 5% (w/v) BSA in PBS. The primary antibodies for detection of IgA (goat-anti-human kappa light chain (#2060-01, Southern Biotech), 1:2,000) or SIRPα-Fc (goat-anti-human IgG Fc-UNLB (#2047-01, Southern Biotech), 1:2,000) were incubated at room temperature for 1 h, before washing with PBS-T. The secondary antibodies (donkey-anti-mouse-AF546 (#A10036, Thermo Fisher Scientific), 1:500; rabbit-anti-goat-AF488 (#A-11078, Thermo Fisher Scientific), 1:500) were combined with Hoechst 33342 (#H1399, Thermo Fisher Scientific) at 10 μg/mL in PBS-TB, and incubated at room temperature in the dark for 45 min. The samples were washed with PBS and imaged using a Leica SP8 confocal microscope (see below).

For the IF-based detection of IgA and SIRPα-Fc on-chip, the cells were seeded in the microfluidic devices as described above and incubated overnight. All incubation steps were performed on the perfusion rocker, unless stated otherwise. For the detection of *in situ* produced proteins, AdV-based gene delivery on-chip was performed following the overnight incubation with cells on-chip, and the chips were cultured for three days as described. For the visualization of binding of purified IgA and SIRPα-Fc, the proteins were added after dilution in the respective cell culture medium to 50 nM, and added on-chip, 50 μl/reservoir. The on-chip cultures were incubated for 1 h. The medium containing the protein standards (controls) was then carefully removed from the reservoirs, and the on-chip cultures were washed with PBS (50 μl/reservoir), 3 × 20 min. The samples were then fixed with 4% (v/v) PFA in PBS at room temperature for 30 min, followed by washing with PBS (3 × 20 min) at room temperature. The samples were blocked with 5% BSA (w/v) in PBS-T for 40 min at room temperature. The primary antibody solutions were prepared as above and added to the on-chip cultures at 50 μl/reservoir. The chips were incubated overnight at 4°C and washed with PBS-T. The secondary antibody solutions were prepared as above and incubated with the on-chip cultures at room temperature for 90 min in the dark, before the final 3 × 10 min washing steps with PBS-T and 1 × 10 min with PBS, and image acquisition with a Leica SP8 confocal microscope.

### 4.10. Confocal microscopy

For confocal microscopy in monolayer cell culture, cells were seeded, incubated and analyzed in ibidi 18-well chamber slides as described above. For the (co-)cultures in the microfluidic devices, images were acquired directly on-chip. A Leica TCS SP8 confocal microscope (Leica Microsystems SP8, Leica, Mannheim, Germany) was used for image acquisition. Hoechst 33342 and CellTrace Violet were excited at 405 nm (detection at 415-465 nm), CellTrace carboxyfluorescein succinimidyl ester (CFSE) and Alexa Fluor 488 were excited at 488 nm (detection at 496-550 nm), CellTrace Yellow at 549 nm (detection at 586-679 nm), Alexa Fluor 569 at 569 nm (detection at 575-620 nm), iRFP at 670 nm (detection at 679-795 nm) and CellTrace Far Red at 633 nm (detection at 641-711 nm). For both monolayer and on-chip imaging, the following objectives were used: HC PL FLUOTAR 10×/0.30 dry, HC PL FLUOTAR 20×/0.50 dry, HCX APO U-V-I 40×/0.75 DRY, and HC PL FLUOTAR 63×/1.2 wet (water immersion). The collected images were processed and quantified using Fiji image analysis software version 1.53v.^95^

### 4.11. Isolation of primary human polymorphonuclear neutrophils (PMN)

The medical scientific research involving human subjects regulated under the Dutch WMO law was approved by Regio Arnhem-Nijmegen medical ethical review board (NL71702.091.20, INVOLVE), and the blood was obtained after the informed consent was given by the volunteers. Fresh blood of healthy donors was obtained by venipuncture of the peripheral vein. The isolation of polymorphonuclear leukocytes (PMN) is later referred to as neutrophil isolation. The neutrophil isolation was performed using the PolymorphPrep density gradient (#1895, Progen, Heidelberg, Germany) according to the manufacturer’s instructions. The erythrocyte contamination was removed by erythrocyte lysis: the washed pellet was resuspended in 45 mL ice-cold milliQ and incubated for 30 s, while gently inverting the tube. Five mL of 10× PBS was immediately added to stop the lysis step, the tube was gently inverted to restore the osmolarity of the solution. The cells were then centrifuged for 5 min at 500 *g* at room temperature, and the pellet was resuspended in cell culture medium containing 10% FCS and 1× PenStrep. The viable cell density was determined by cell counting with trypan blue staining, and adjusted to the desired amount with the appropriate cell culture medium. 10 pg/mL of Granulocyte-Macrophage Colony-Stimulating Factor (GM-CSF, #130-093-862, Miltenyi Biotec, Bergisch Gladbach, Germany) was added immediately before the start of the assay to activate the neutrophils before the ADCC assay.

### 4.12. Macrophage differentiation and polarization

Monocytes were isolated from freshly isolated or cryopreserved peripheral blood mononuclear cells (PBMCs) and subsequently differentiated and polarized into M2-like macrophages as reported before.^33^

### 4.13. Image-based ADCC assay with primary human neutrophils in monolayer cultures

Target (cancer) cells were stained with 2.5 µM CellTrace Far Red (Thermo Fisher Scientific) according to the manufacturer’s instructions, and seeded in a flat bottom 96-well plate (#3596, Corning) at a density of 2 × 10^3^ cells/well. Each sample was prepared as at least in duplicates. Following an overnight incubation, the medium was replaced with medium containing a 1:1,000 dilution of CellTox Green viability dye (#G8741, Promega, Mannheim, Germany) and recombinant IgA and/or SIRPα-Fc proteins was added. Freshly isolated neutrophils were added to the samples at a density of 3 × 10^5^ cells/well for HCT116 and 2.5 × 10^5^ cells/well for MDA-MB-468 cells, considering the doubling time of each cell line, to achieve an effector-to-target cells (neutrophil-to-cancer cell) ratio (ET ratio) of 50:1. The final volume of each well was 200 µL. The plate was centrifuged at 500 *g* for 3 min to let the neutrophils settle at the bottom of the well. The plate was placed in the Incucyte ZOOM live-cell analysis system (Sartorius, Göttingen, Germany) with 4× objective, and the images of phase contrast and red and green fluorescence channels were acquired in one-hour intervals. The ADCC was quantified using the Incucyte Zoom software package (Sartorius) as follows: six representative images were selected from each plate as training images to create a processing definition in “Basic Analyzer”. The parameters were adjusted to achieve the optimal masking of the cells in the green, red and overlap channels. The established processing definition was then used to process all the images. The cancer cell mask was obtained from the red channel (CellTrace Far Red), while the dead cell area was taken from the green channel (CellTox Green). Areas with co-localization of green and red signals indicated dead cancer cells. To quantify the ADCC, the percentage of overlapping area from the total red area was calculated. The value at t=0 (the percentage of the area overlap at the start of the assay) was subtracted from the values of each of the time points as a background value of the tumor cell death prior to start of the assay. The background-subtracted values for the period of eight hours were plotted using the GraphPad Prism software v.9.5.1.

### 4.14. ADCC assay on-chip

Target (cancer) cells were initially stained with 1 µM CellTrace Red (Thermo Fisher Scientific) according to the manufacturer’s instructions, and seeded on-chip in collagen as described above at a density of 1.2 × 10^4^ cells/chip (loading volume: 6 µL). In the ADCC assay upon AdV transduction, virus-based delivery was performed as described above. One day before the start of the assay, CellTox Green viability dye was added in the medium on-chip at 1:2,000 dilution. Three days post transduction, the images of representative areas of the on-chip culture were captured using a fluorescent microscope (Evos, Life Technologies, Thermo Fisher Scientific) in the red and green channel (capturing the CellTox Green and CellTrace Far Red signal for the dead cells and tumor cells, respectively). The images provided the background values of the tumor cell death per each chip prior to addition of effector cells. The ADCC assay was started by addition of effector cells and recombinant IgA and/or SIRPα-Fc as controls. IgA and SIRPα-Fc proteins were added on-chip in cell culture medium, 25 µL/reservoir, containing CellTox Green viability dye at a final dilution of 1:2,000. Isolated primary human neutrophils were resuspended and added to the upper side channel reservoirs at 1 × 10^7^ cells/mL in 100 µL. N-formyl-methionyl-leucyl-phenylalanine (fMLP) was prepared at 2× of the desired final concentration of 100 nM and was added to the lower channel to create a chemoattractant gradient for the neutrophils and promote the neutrophil invasion into the tissue compartment at the beginning of the assay. Chips were incubated overnight on a perfusion rocker. Following the incubation, images of two representative areas per chip were acquired with a fluorescent microscope as above, and Fiji ImageJ software was used for image analysis. The ADCC was quantified as follows: CellTrace Far Red signal was thresholded to create a cancer cell mask, and the area of the mask was measured as the total tumor cell area. The CellTox Green signal was thresholded for the dead cell mask. The overlapping area of CellTrace Red mask and CellTox Green mask was measured and taken to represent dead cancer cells. The percentage of dead cancer cells was calculated by taking the ratio between ‘dead cancer cell’ and ‘total cancer cell’ area. The background value of cancer cell death per each chip was subtracted from each of the respective endpoint values, and the average background-corrected normalized cancer cell death value per chip was calculated. The ADCC assay for investigation of the treatment specificity was performed as described, with cancer cells and primary fibroblasts labeled with CellTrace Far Red and CellTrace Violet, respectively, and seeded in collagen as described above at a 1:1 ratio to the final cell density of 5×10^3^ cells/chip. Following the AdV transduction and addition of the effector cells as described above, the chips were imaged with a Leica SP8 confocal microscope using the HC PL FLUOTAR 10×/0.30 dry objective. The images were processed as described above, with the mask and the overlapping area representing dead cells calculated per cell type.

### 4.15. ADCP assay on-chip

Target (cancer) cells were initially stained with 0.7 µM pHrodo Red pH-sensitive dye (Thermo Fisher Scientific). M2-polarized macrophages were labeled with 0.75 µM CellTrace Violet cytoplasmic dye. The cells were combined in an E:T ratio of 1:1.25 in 3 mg/mL collagen, and seeded on-chip at the density of 2×10^4^ cells/mL in a volume of 6 μL. The collagen preparation and cell seeding were described above. Following the gelation of collagen, the medium containing AdV-adapter complexes or recombinant proteins as controls were added to the side channels. The HCT116 target cells were thereby transduced with adenoviral particles encoding eGFP (vehicle control), or anti-EpCAM IgA or anti-EpCAM dimeric IgA at an MOI of 5 i.v.p./cell, retargeted with the EGFR-binding E01 DARPin-based adapter. The MDA-MB-468 target cells were transduced with adenoviral particles encoding eGFP (vehicle control), or anti-EGFR IgA, anti-EGFR dimeric IgA, SIRPα-Fc or a combination thereof, at an MOI of 5 i.v.p./cell, and retargeting occurred with an EpCAM-binding Ac2 DARPin-based adapter. The chips were incubated overnight on a perfusion rocker as described, the confocal images of at least two representative areas and planes per chip were acquired using a Leica SP8 confocal microscope. All images were acquired using the HCPL FLUOTAR 20×/0.5 dry objective and analyzed with the Fiji ImageJ software. The phagocytic index was determined based on a modified algorithm for ADCP quantification on-chip we reported earlier^33^. As at least two areas per chip were analyzed, the resulting phagocytic index per chip was calculated as an average of the indexes of the respective chip.

### 4.16 Animal experiments

SCID mice (Charles River, 11 weeks old, female) and C57BL/6 mice (Janvier, 13 weeks old, female) were housed in a Biosafety Level 2 (BLS2) facility within individually ventilated cages (IVCs), 3-4 animals per cage. All mice experiments were performed in accordance with the Swiss animal protection law and with approval of the Cantonal Veterinary Office (Zurich, Switzerland).

For the tumor treatment models the mice were bred at Janvier Labs (France) and transported to the animal facility of the Utrecht University (GDL) at least 1 week prior to each experiment. Food and water were provided ad libitum and mice were housed in groups under a 12:12 light–dark cycle. Mice were sacrificed by cervical dislocation. Female human FcαRI (CD89) transgenic^42^ or non-transgenic littermates from C57BL/6JRj (B16F10_luc2-EGFr) or CB17-SCID (HCT116 and MDA-MB-468) strains were used. Mice in experimental groups were randomized based on genotype and tumor volume and researchers were single-blinded. All mouse experimental procedures were approved by the institute’s animal ethics committee and by the Dutch Central Authority for Scientific Procedures on Animals (CCD, AVD11500202115442) On the day of tumor cell injection, HCT116, MDA-MB-468, and B16F10-EGFR cells were harvested upon reaching ∼80% confluency, washed and resuspended in PBS to the desired density. The following cell densities were used: HCT116/MDA-MB-468 at 2.5 million/mouse, and B16F10-EGFR at 0.75 million/mouse. Cell density was adjusted by mixing 130 µL ice-cold Growth Factor Reduced Matrigel (Ref. 356231, Corning) with 870 µL cell suspension per mL. For each mouse, 100 µL was prepared. Before injection, the mixtures were transferred to 1 mL syringes with 26G needles and maintained on ice until use. The cell suspension was injected subcutaneously in a right flank after mice were put under anesthesia using isoflurane vaporizer. Tweezers were used to gently pinch the skin around the needle upon withdrawal to prevent reflux.

Tumor outgrowth was measured after the tumors became palpable using an electronic caliper three times a week. Tumor volume for non-transgenic mice was calculated as V = 0.5 * L * W^2^ (L = length, mm; W = width, mm). For *in vivo* studies performed with FcαRI transgenic mice, which were done at a different site, an alternative method was utilized: L × H(eight) × W. Tumor skin was monitored for inflammation and breakage once tumors exceeded 500 mm^3^, and humane endpoints were followed. No signs of treatment-related toxicity or weight loss in the animals were observed throughout the duration of treatment.

When the tumors reached a volume of approximately 150 mm^3^ (between 50-300 mm^3^), the mice were intratumorally injected with the retargeted AdV or PBS control as follows: AdV, adapter, and shield preparation solutions, prepared as reported before^44^, were thawed on ice. Briefly, the shield protein (anti-hexon binding scFv, trimerized) was expressed by baculovirus-transduced Sf9 culture. The secreted protein was isolated by protein L affinity chromatography and further purified by size exclusion chromatography. Purity was validated by SDS-PAGE and mass spectrometry and mono-dispersity was confirmed by size exclusion HPLC. Shield protein used in the *in vivo* study was free of potential endotoxin contaminations. For each tumor, 50 µL of the virus mix containing 11 × 10^9^ shielded viral particles (o.v.p.) was prepared for intratumoral injection per tumor. Complexes were prepared by pre-diluting adapters to 5.0 nM in PBS, pipetting virus solutions into tubes, pipetting adapter into the virus solutions and mixing immediately, followed by a 40-min incubation at room temperature. Shield solutions were then added, mixed, and incubated for an additional 30-40 min. The final solution was adjusted with PBS and kept at room temperature before injection.

The viral solution was injected using 1 mL syringes with 27-30G needles, aiming for the middle of the tumor. Mice were anesthetized as previously described. The tumor skin and underlying tumor were lifted with tweezers, and the needle was inserted through the skin approximately 1 cm below the tumor. Proper needle placement was confirmed by gently moving the needle and ensuring the tumor moved accordingly before injection. Plasma samples were collected during the study by drawing blood via a cheek pouch puncture starting 7 days after the adenovirus injection which was repeated every 7 days or when mice reached humane endpoint. For the confirmation of delivery study, animals were euthanized 4 days after the virus injection using carbon dioxide. For the confirmation of activity study in FcαRI transgenic mice, mice were sacrificed at the end of the study or when reaching humane endpoints. Tumors were carefully resected using a scalpel and scissors and processed. For RNA isolation, tumor material was washed in ice-cold PBS followed by subsequent flash-freezing for RNA isolation and storage at -80°C. For analysis by IF, tumors were washed in PBS for 1 min and fixed overnight in 4% PFA in PBS at 4°C. Tumors were placed in 30% (w/v) sucrose in PBS for 2 days at 4°C. Tumors (> ∼5 mm length) were cut in half, each half embedded separately with either inner or outer side towards the bottom of the mold, to section the respective region of the tumor. Tissues were then covered with O.C.T. Tissuetek and placed in 10×15 mm peel-away mold, after which they were frozen by immersion in dry ice-cooled 2-methylbutane at -40°C for 2 min. Tissue blocks were sectioned using a cryostat into sections of 8 µm for subsequent analysis by IF. Tissue sections were stored at -80°C until use.

### 4.17. Immunofluorescence analysis of tumor sections

Slides with tissue sections were thawed to room temperature and fixed with 4% (v/v) PFA in PBS on the glass for 10 min at RT. Each section was outlined using a liquid-blocking sample pen. Slides were washed 3 × 5 min with PBS. B16F10-EGFR tumor sections were additionally bleached with 10% H_2_O_2_ in PBS for 1 h at RT, after which they were washed 3 × 5 min with PBS. Sections were treated with the primary antibody in PBS with 0.05% Tween-20 and 0.5% BSA (PBS-TB) for 1 h at RT. Unlabeled primary antibodies used were anti-EGFR IgA (0.3 mg/ml; 1:300; for HCT116 and B16F10) and SIRPα-Fc (0.3 mg/ml; 1:300) for detection of EGFR or CD47, respectively. For detection of the locally produced IgA and SIRPα-Fc fusion protein, goat anti-Ckappa light chain (1:1,000) and goat anti-human Fc gamma (1:1,000 dilution) were used, respectively. Slides were washed 3 × 5 min with PBS and incubated with the secondary antibody in PBS-TB for 45 min at RT in the dark. Secondary or directly labeled antibodies used were donkey-anti-goat-AF488 (1:600), rat anti-CD45-AF594 (1:600), rat anti-Ly6G-AF647 (1:600) or mouse anti-EpCAM-AF594 (1:600) for detection of primary antibodies against CD45-positive immune cells, Ly6G-positive immune cells or EpCAM, respectively. The samples were counterstained with 300 nM DAPI (Thermo Fisher Scientific) for 5 min at RT. After final washing, sections were mounted with ProLong Gold antifade mounting agent (Thermo Fisher Scientific) and air-dried overnight. Confocal microscopy analysis was performed as described above. Full section images were acquired using a TileScan function. The images were processed and analyzed as described above.

### 4.18. Measurement of titers in plasma

Nunc MaxiSorp flat-bottom 96-well plates (Thermo Fisher Scientific) were coated with 50 μl/well goat-anti-human kappa (Southern Biotech; #2060-01 1:2000) for the IgA ELISA or with rabbit-anti-human IgG (Jackson art#309-006-008; 0.5 μg/ml) in PBS for the SIRPα-Fc ELISA and incubated overnight at 4°C. The next day, the coating solution was aspirated and the plate was washed 3x with PBS-T. The wells were blocked with 100 μl/well of 0.5% (w/v) BSA in PBS for 1 h at room temperature, followed by 3 × washing with PBS-T. The titrated standards for estimating plasma concentrations: purified anti-GD2 IgA3.0, anti-GD2 IgG1, or SIRPα-Fc, blanks, or the sample dilution series (plasma samples diluted in 0.01% (v/v) Tween-20 + 0.5% (w/v) BSA in PBS (PBS-TB)) were added, 50 μl/well. The samples were incubated for 2 h at room temperature, followed by a 3 × washing with PBS-T. For the IgA detection with goat-anti-human IgA HRP; (Southern Biotech; #2050-05, 1:2000) was used, 50 μl/well was added and left to incubate for 1 h at room temperature. For SIRPα-Fc detection, Goat-anti-human IgG HRP; (Jackson art# 109-035-098, 1:2,000), 50 μl/well was added and left to incubate for 1 h at room temperature. The plate was washed as above and ABTS substrate (Bioquest, cat# 11001), 50 μl/well was added. The signal was measured with a Microplate Reader (SpectraMax M3, Molecular Devices) by recording absorbance at 405 nm. Once the optimal signal was reached, The data were analyzed using GraphPad Prism Software v.9.5.1 for Windows.

### 4.19. Gene expression analysis of tumor tissues<colcnt=6>

Tumor samples were dissociated with the VDI 12 homogenizer (VWR®) in 3 ml of TRI Reagent (#T9424, Merck) and RNA was extracted using chloroform. The RNA was purified by washing with isopropanol and ice-cold 75% ethanol before resuspending it in MilliQ. RNA yield and purity was assessed using NanoDrop (ThermoFisher Scientific), and 1000 ng of RNA was used for cDNA synthesis. After incubation with random primers (#C1181, Promega) and deoxynucleotides (#N0447L, New England BioLabs) at 65°C for 5 min, a mix of RNAse inhibitor (#M0307S, New England BioLabs), dithiothreitol (DTT; #Y00147, Thermo Fisher Scientific), 5× first strand buffer (Thermo Fisher Scientific) and SuperScript II Reverse Transcriptase (#18064014, Thermo Fisher Scientific) was added according to the manufacturer’s instructions. The cDNA was generated by incubation of the samples for 10 min at 20°C, followed by 50 min at 42°C, and 3 min at 59°C. cDNA was stored at -20 °C until further analysis using qPCR. The following primer pairs (forward (F) and reverse (R), as indicated) were used for amplification of the murine gene transcripts: mIFN-γ: (F) 5’-CAGCAACAGCAAGGCGAAAAAGG-3’, (R) 5’-TTTCCGCTTCCTGAGGCTGGAT-3’; mTNF-α: (F) 5’-GGTGCCTATGTCTCAGCCTCTT-3’, (R) 5’-GCCATAGAACTGATGAGAGGGAG-3’; mIL-1β: (F) 5’-TGGACCTTCCAGGATGAGGACA-3’, (R) 5’-GTTCATCTCGGAGCCTGTAGTG-3’; mIL-6: (F) 5’-TACCACTTCACAAGTCGGAGGC-3’, (R) 5’-CTGCAAGTGCATCATCGTTGTTC-3’; mIFN-β: (F) 5’-GCCTTTGCCATCCAAGAGATGC-3’, (R) 5’-ACACTGTCTGCTGGTGGAGTTC-3’; mHPRT1 (reference gene): (F) 5’-GGACTGATTATGGACAGGACTG-3’, (R) 5’-GCTCTTCAGTCTGATAAAATCTAC-3’; mTBP-1 (reference gene): (F) 5’-CTACCGTGAATCTTGGCTGTAAAC-3’, (R) 5’-AATCAACGCAGTTGTCCGTGGC-3’. qPCR was performed in a CFX Connect Real-Time System (Bio-rad) by mixing cDNA with primer pairs at final concentration of 400 nM, iQ SYBR Green Supermix (#1708882, Bio-rad), and MilliQ, at final volume of 12.5 μl. Target was compared with the geometric mean of reference genes CT values for each experimental condition (ΔCT).

### 4.20. Statistical analysis

To reliably analyze pooled data from multiple ADCC or ADCP experiments and minimize effects of donor variation, each of the values within each biological experiments was normalized for the total sum of values from that experiment, resulting in a normalized ADCC or ADCP index. A p-value of <0.05 was considered statistically significant. For comparisons between two groups, two-tailed unpaired t-tests were performed, assuming normally distributed data; for the comparison of the parallel kinetic measurements (ADCC in monolayer culture), a two-tailed paired t-test was performed between the test and control values at each of the time points. For comparisons between multiple groups, one-way ANOVAs with Dunnett’s correction for multiple comparisons were performed. For the comparison of neutrophil-mediated killing of cancer cells and fibroblasts (treatment specificity analysis), a two-tailed unpaired t-test was performed between the cancer cells and fibroblasts, with two-stage step-up (Benjamini, Krieger, and Yekutieli) correction for multiple comparisons. Statistical analyses of ADCC and ADCP assays were performed using GraphPad Prism v1.53v. Statistical analysis of treatment-induced changes in tumor growth was done using a mixed model approach, with a Geisser-Greenhouse correction. Correction for multiple comparison was done with Dunnett’s multiple comparisons test, whereby each treatment was compared with vehicle control across timepoints. Adjusted p values < 0.05 were considered significant. Survival curves were analyzed using the Kaplan-Meier method and Log-Rank test, with a Bonferroni correction applied for multiple testing. Adjusted p values < 0.05 were considered significant. Results from mice experiment were analyzed with GraphPad Prism 10.1.2.

## Data availability statement

All data are available in the main text, figures, tables or supplemental information.

## Acknowledgements

M. Chernyavska was supported by an internal funding from Radboudumc. The authors thank the Radboudumc Technology Center Microscopy for use of their microscopy facilities.

## Author contributions

M. Chernyavska: Conceptualization, and study design, Methodology, Investigation, Formal Analysis, Writing – Original Draft Preparation, Writing – Review & Editing; K.P. Hartmann: Methodology, Investigation, Writing – Review & Editing; J.H.M. Jansen, Methodology, Investigation, Formal Analysis, Writing – Review & Editing; N. Baumann: Investigation, Writing – Review & Editing; J. Kolibius: Methodology, Investigation, Writing – Review & Editing; D. Brücher: Methodology, Investigation, Writing – Review & Editing; T. Kristoforus: Investigation, Writing – Review & Editing; R.H.W. Peters: Investigation, Writing – Review & Editing; L. Huijs: Investigation, Writing – Review & Editing; D. Laarveld: Investigation, Writing – Review & Editing; F. Weiss: Investigation, Writing – Review & Editing; ; R. Burger: Investigation, Writing – Review & Editing; M. Lustig: Investigation, Writing – Review & Editing; N. Gimenez de Assis: Investigation, Writing – Review & Editing; M. Schmid: Conceptualization, Resources, Methodology, Writing – Review & Editing; J.H.W. Leusen: Conceptualization, Methodology, Supervision, Writing – Review & Editing; T. Valerius: Conceptualization, Methodology, Supervision, Resources, Writing – Review & Editing; A. Plückthun: Conceptualization, Methodology, Supervision, Writing – Review & Editing; W.P.R. Verdurmen Conceptualization, Methodology, Supervision, Writing – Original Draft Preparation, Writing – Review & Editing.

## Declaration of interests

The authors declare no competing interests

## Notes

### Competing Interest Statement

The authors have declared no competing interest.

