## Supplementary Materials for "Retargeted adenoviruses for local IgA and CD47 blocker production as a novel cancer therapy"

Mariya Chernyavska et al.

### Supplementary material

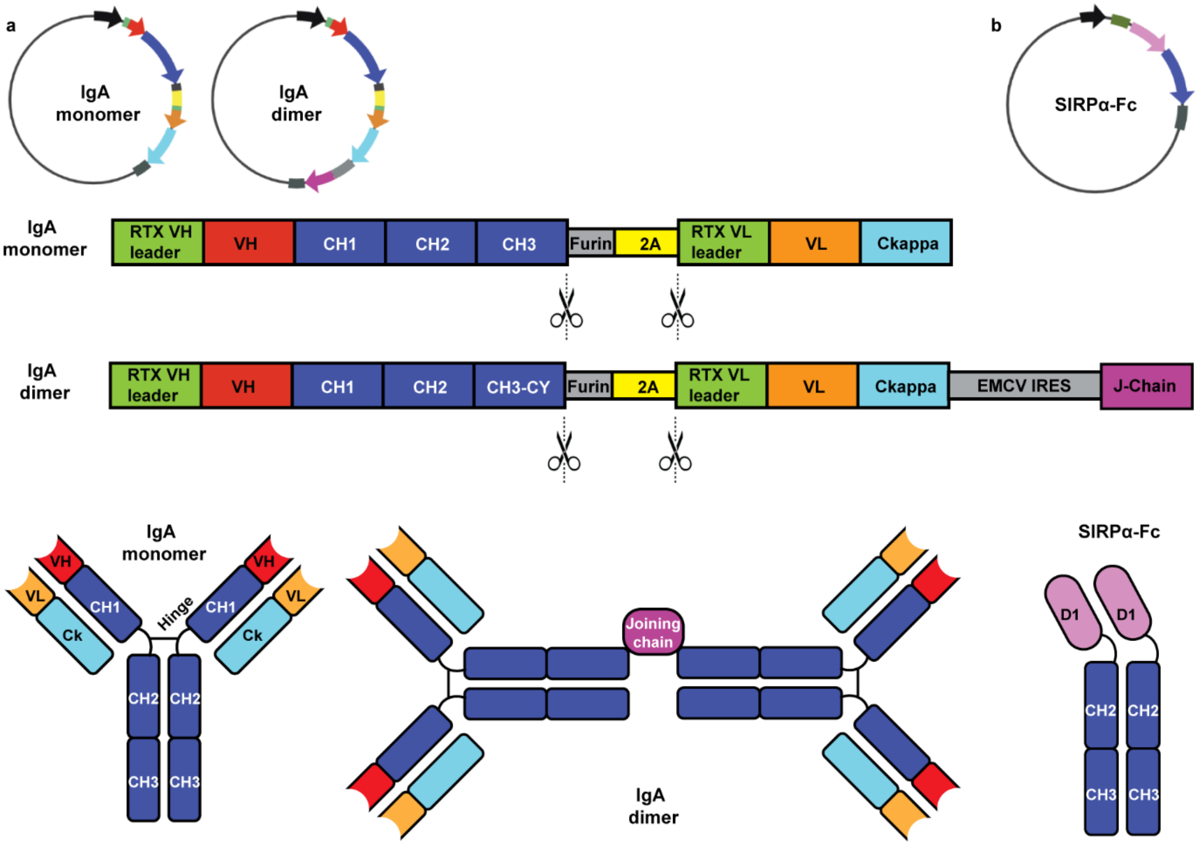


**Supplementary Figure S1. Schematic representation of the engineered pShuttle expression vectors for expression of IgA (a) and SIRPα-Fc (b). (a)** Schematic representation of the IgA molecules and the corresponding vector scheme maps are shown. Black: promoter (CMV); green: leader sequence from rituximab; red, orange: VH and VL variable domains of the heavy and light chain of IgA, respectively; gray: furin recognition sequence, followed by a P2A or T2A "self-cleaving" (ribosome skipping) sequence (yellow); the cleavage site is indicated schematically; blue and cyan: heavy chain and light chain of IgA with the respective CH1-CH3 heavy and VL light chain domains. Dimeric IgA was encoded in the same vector format with a modified CH3 domain (introduced C-terminal Cys-Tyr (CY) residues), with the joining (J-) chain (magenta) encoded under EMCV IRES sequence (light gray). **(b)** Schematic representation of the SIRPα-Fc molecule and the respective expression vector map. Black: EF-1α promoter; green: SIRPα leader sequence; pink: Consensus variant (CV) of D1 domain of SIRPα engineered with increased affinity to human CD47 and cross-reactivity with murine CD47; blue: Fc-ablated, AAA-mutant variant of the IgG1 heavy chain CH2-CH3 domains. RTX, rituximab.

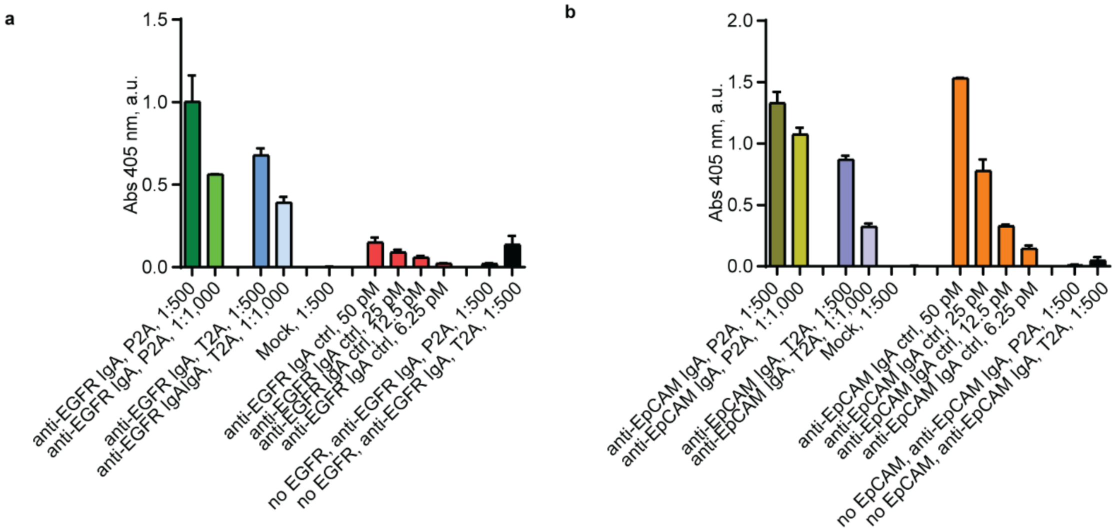


**Supplementary Figure S2. Expression of IgA antibodies from P2A and T2A vector variants compared in ELISA**. **(a)** ELISA-based detection of anti-EGFR IgA in supernatants (at 1:500 and 1:1,000 dilution) of CHO-S cells transfected with the IgA-encoding constructs, containing either P2A or T2A “self-cleaving” (ribosome-skipping) sequences. Mock-transfected (eGFP construct) CHO-S supernatant was used as a negative control. Recombinant anti-EGFR IgA produced by co-expression of heavy and light chains in trans was used as a positive control. To evaluate non-specific binding, a control without coated EGFR domains was included. **(b)** Same as (a), for anti-EpCAM IgA. Representative data from three independent experiments is shown (mean ± range from two technical replicates indicated).

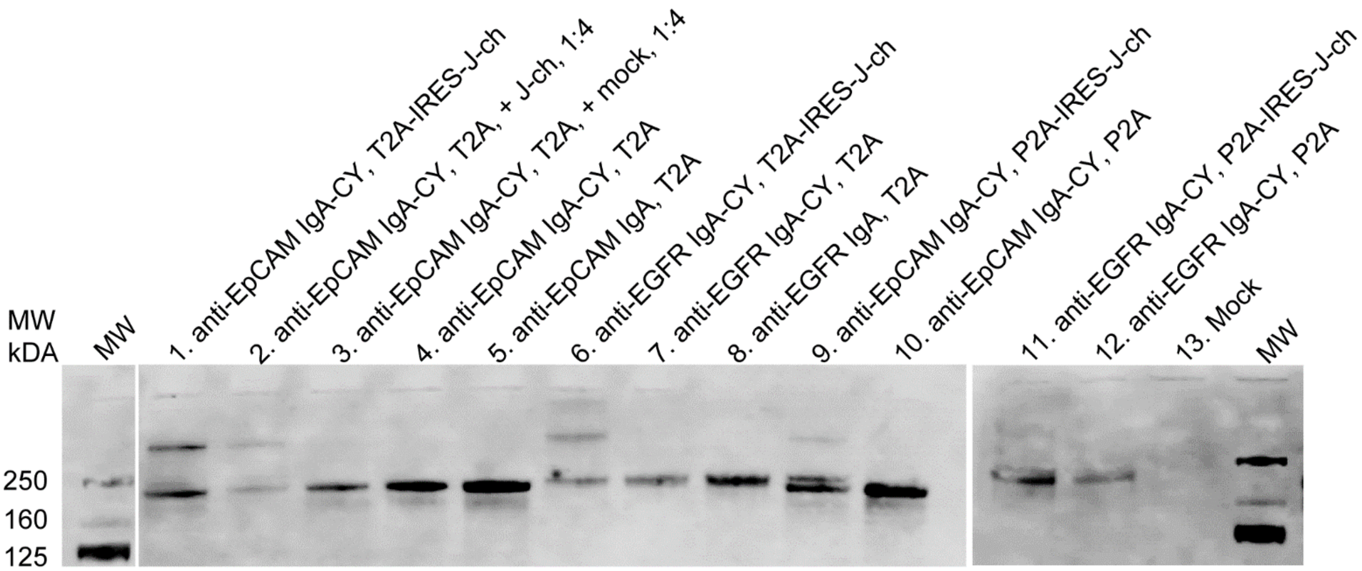


**Supplementary Figure S3. Western blot analysis of dimeric IgA expressed from various expression vectors.** Supernatants of CHO-S cells transfected with the anti-EGFR or anti-EpCAM IgA expression constructs containing T2A or P2A "self-cleaving" (ribosome skipping) sequences were compared in western blot under non-reducing conditions, by detecting IgA heavy chain (HC). Bicistronic constructs encoding monomeric IgA (T2A variants) were used as controls yielding a band corresponding to a monomeric IgA (lanes 5 and 8); the same constructs containing C-terminal Cys-Tyr (CY) residues, which are reintroduced as they are required for dimer formation, were used as controls to validate successful antibody production (lanes 4 and 7 for T2A variants, lanes 10 and 12 for P2A variants). Tricistronic vectors where the J-chain was encoded under an IRES resulted in expression of the dimeric IgA fraction detected at the height of ~350 kDa (lanes 1, 6 for T2A-variants and lanes 9, 11 for P2A variants), with the T2A variants yielding higher proportion of the dimer compared to the monomer fraction (~350 kDa band vs ~170 kDa band). To investigate if supplying J-chain in a higher molar ratio (1:4 IgA construct: J-chain construct) would result in a higher dimer proportion, CHO-S cells were co-transfected with the vector encoding the IgA monomer with the re-introduced CY sequence, together with the vector encoding only the J-chain (lane 2), which resulted in a similar ratio of dimer:monomer as for the tricistronic vector (lane 1) but overall lower yield. To normalize for differences due to co-transfection, we included the control where CHO-S were co-transfected with the same IgA construct as in the lane 2, but with a mock (GFP) construct instead of the J-chain, in the same molar ratios (lane 3). Finally, a mock-transfected sample (GFP) was included as a negative control (lane 13). The image is representative of at least three independent experiments.

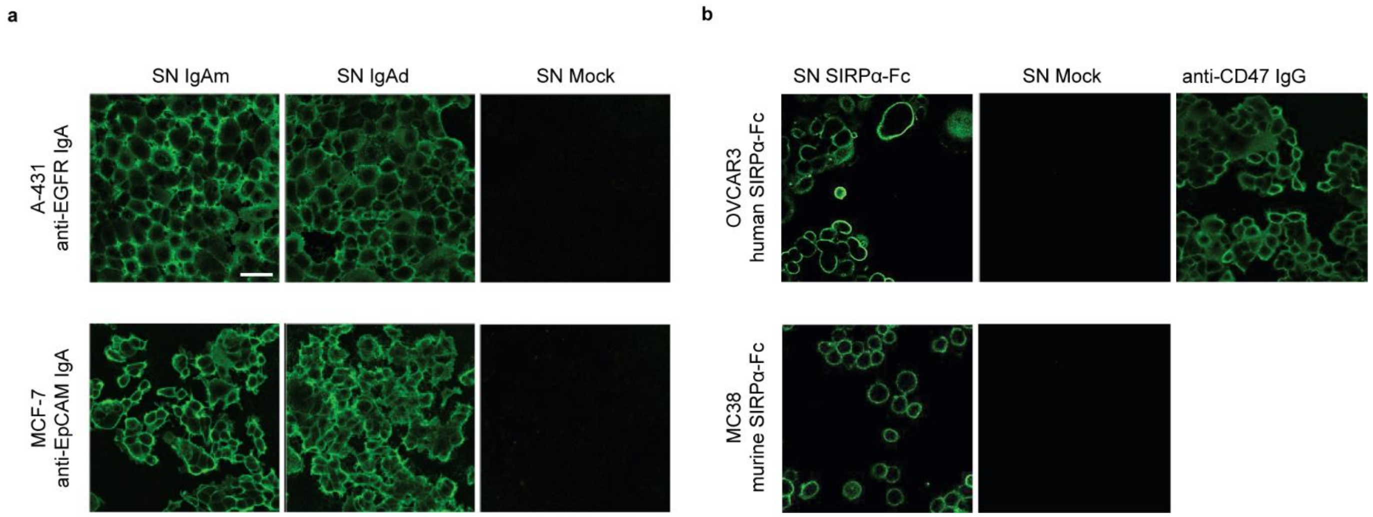


**Supplementary Figure S4. Immunofluorescence-based confirmation of IgA antibody and SIRPα-Fc production in CHO-S cells through target binding in cancer cell lines.** Upon transfection of CHO-S cells with the respective expression vectors, the supernatants (SN) containing the molecule of interest were incubated with the tumor cells expressing the respective target surface receptor, and the binding of the target molecule to the cell surface was visualized by indirect immunofluorescence. **(a)** Supernatants from CHO-S cells transfected with the expression constructs for anti-EGFR or anti-EpCAM IgA monomer (IgAm) or dimer (IgAd), as well as mock-transfected CHO-S supernatants (GFP expression construct) were analyzed by incubating the SN with EGFR-overexpressing A-431 cells, or EpCAM-overexpressing MCF-7 cells, respectively. Note that the dimer-encoding vector produces a combination of monomer and dimer and that we stain for IgA irrespective of its multimerization state. **(b)** Supernatants from CHO-S cells transfected with the SIRPα-Fc-expressing construct or GFP-expressing construct (SN Mock) were analyzed by incubating the SN with OVCAR3 cells (expressing human CD47) or MC38 cells (expressing murine CD47). Positive control: anti-human CD47 IgG. The representative images from three independent experiments are shown. Scale bar: 100 µm.

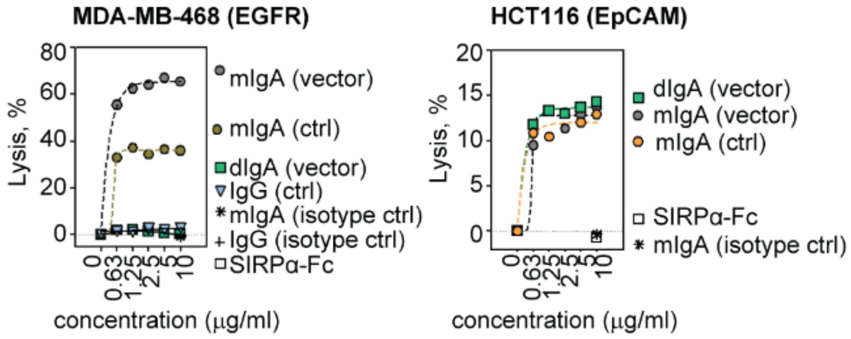


**Supplementary Figure S5. Assessment of the ability of the purified IgA antibodies to induce antibody-dependent cell-mediated cytotoxicity (ADCC) of tumor cells.** MDA-MB-468 and HCT116 cells were used as target cells for anti-EGFR and anti-EpCAM IgA-mediated killing by primary human neutrophils, respectively. IgA/IgG ctrl refers to pure proteins produced using conventional approaches as opposed to purified from supernatants of adenoviral vector-transduced cells. For MDA-MB-468 cells, IgG was used as a negative control for the assay. Of note, for the purified dimeric mix of anti-EGFR IgA, we observed aggregation and a complete lack of cell-killing activity. N=3, a representative experiment is shown.

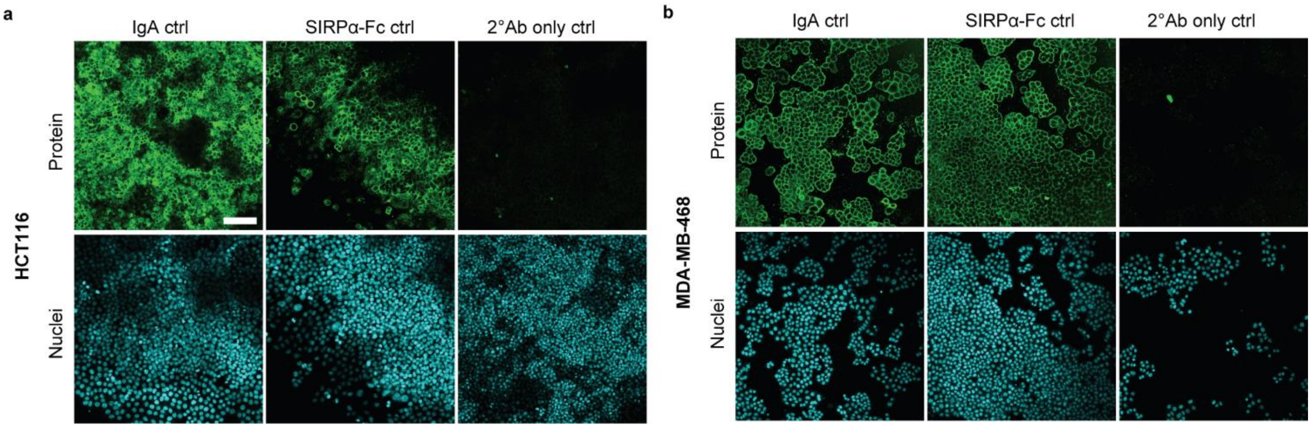


**Supplementary Figure S6. Immunofluorescence-based detection of IgA and SIRPα-Fc recombinant controls in HCT116 and MDA-MB-468 cell lines.** Supplementary panels for Figure 2. **(a)** Binding of anti-EpCAM IgA and SIRPα-Fc recombinant proteins at 20 nM was detected by using goat-anti-human kappa light chain and goat anti-human IgG Fc as primary antibodies, respectively. Staining with a secondary rabbit anti-goat antibody only (2º Ab only ctrl) was performed as a negative control for both IgA and SIRPα-Fc. **(b)** Same as (a), for anti-EGFR IgA and SIRPα-Fc recombinant proteins in MDA-MB-468 cells. Scale bar: 100 µm.

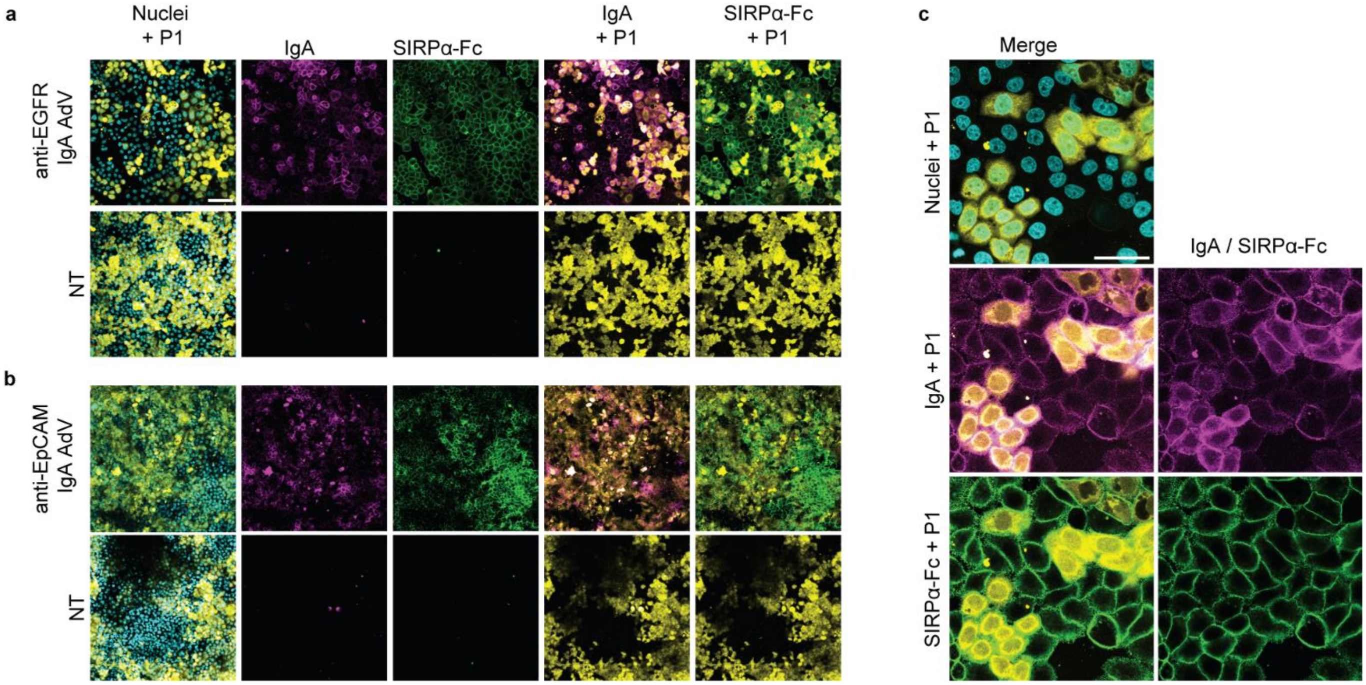


**Supplementary Figure S7. IgA antibody and SIRPα-Fc genes are co-delivered with retargeted adenoviral vectors and produced proteins bind in an autocrine and paracrine fashion to tumor cells.** Anti-EGFR IgA and anti-EpCAM IgA were produced in MDA-MB-468 **(a, c)** and HCT116 cells **(b)** upon retargeting of the AdV vector to either EpCAM or EGFR. Cells were labeled with cell-permeant non-fluorescent CellTrace dyes which react with amino groups and become fluorescent. Yellow, population one of the tumor cells (P1, labeled cell population) transduced with AdV in the respective samples; cyan, nuclei (Hoechst, both populations); magenta, IgA (detected by goat anti-human kappa light chain); green, SIRPα-Fc (detected by goat anti-human IgG Fc). The merge images (cyan and yellow) show the overlay between the nuclei (both populations) and the transduced cells (P1). All non-yellow areas indicate staining on non-transduced cells in a paracrine fashion. Scale bar: 100 µm. **(c)** A close-up of MDA-MB-468 cells producing IgA and SIRPα-Fc in a paracrine fashion. Cells were not permeabilized before staining. NT, non-treated. Scale bar: 50 µm. Positive controls (purified recombinant IgA, SIRPα-Fc proteins) and secondary Ab only control images are shown in Figure S8. MOI: 20 i.v.p./cell (each virus). Representative images from three independent experiments are shown.

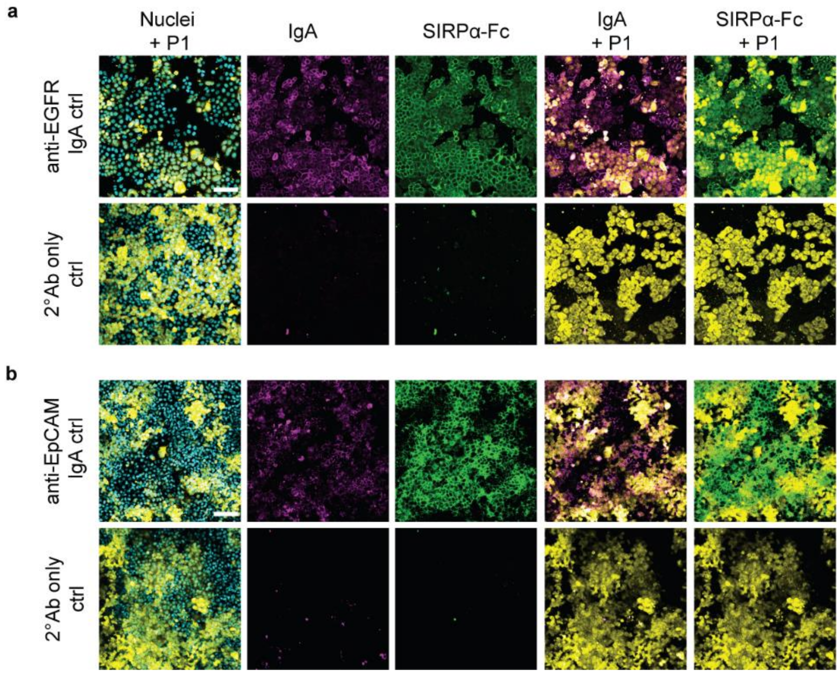


**Supplementary Figure S8. IgA and SIRPα-Fc co-delivered to tumor cells** (additional controls for Figure S7). **(a)** MDA-MB-468 cells were seeded into the same well in two populations, one population labeled with CellTrace Yellow (P1), and one non-labeled. Nuclei of both populations were stained with Hoechst (cyan). As a positive control for co-delivery of IgA and SIRPα-Fc, the recombinant proteins at 20 nM each were added to the tumor cells and detected simultaneously in indirect immunofluorescence. Magenta: anti-EGFR IgA; green: SIRPα-Fc. Similar to Figure S7, the overlay images of IgA or SIRPα-Fc with the labeled tumor cell population are shown. **(b)** Same as (a), for HCT116 cells, and anti-EpCAM IgA. Scale bar: 100 μm.

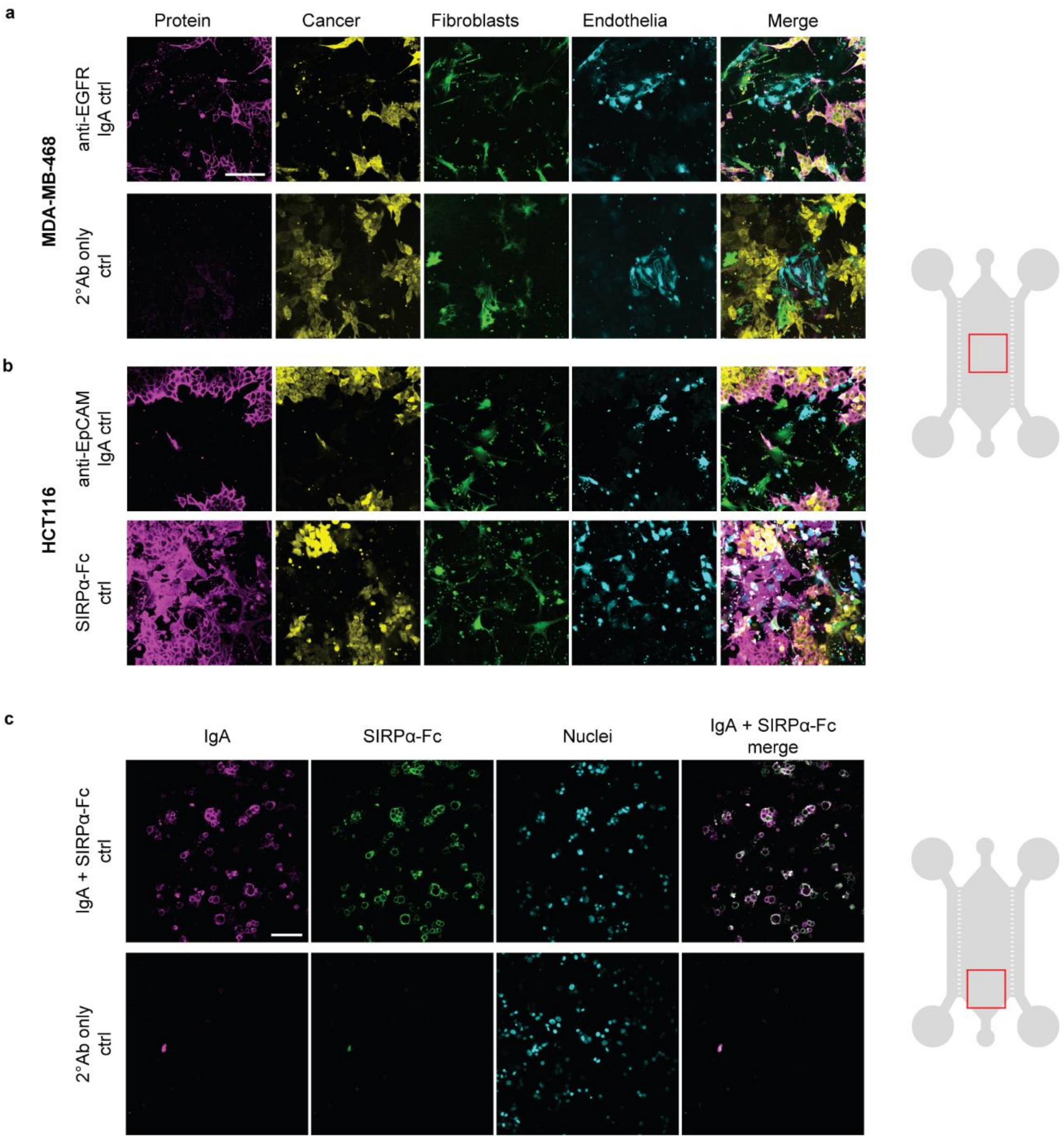


**Supplementary Figure S9. IgA and SIRPα-Fc co-delivered to the tumor microenvironment on-chip.** Supplementary panels for Figure 4. **(a)** Anti-EGFR IgA (magenta) was added at 20 nM as a recombinant protein control; secondary antibody only (2ºAb only ctrl) is shown. As in Figure 4, the schematic image represents the tumor-on-a-chip with an approximate image area indicated. Yellow (CellTrace Far Red-stained): MDA-MB-468 cells; green (Cell Trace yellow-stained): fibroblasts; cyan (CellTrace Violet-stained): endothelial cells; overlay images demonstrate the specificity of IgA binding on the surface of the cancer cells. **(b)** Recombinant controls for anti-EpCAM IgA and SIRPα-Fc (magenta) in HCT116 tumor-on-a-chip model, similar to (a). Anti-EpCAM IgA showed specificity for cancer cells, whereas SIRPα-Fc was detected on the surface of all cell types. (**c**) Positive control and secondary antibody only control for Figure 4c. Magenta: recombinant anti-EGFR IgA (20 nM); green: recombinant SIRPα-Fc (20 nM); cyan: nuclei. Similar to (a) and (b), the tumor-on-a-chip schematic image shows the approximate area imaged. Representative images are shown from at least two chips in at least two independent experiments. Scale bar: 100 μm.

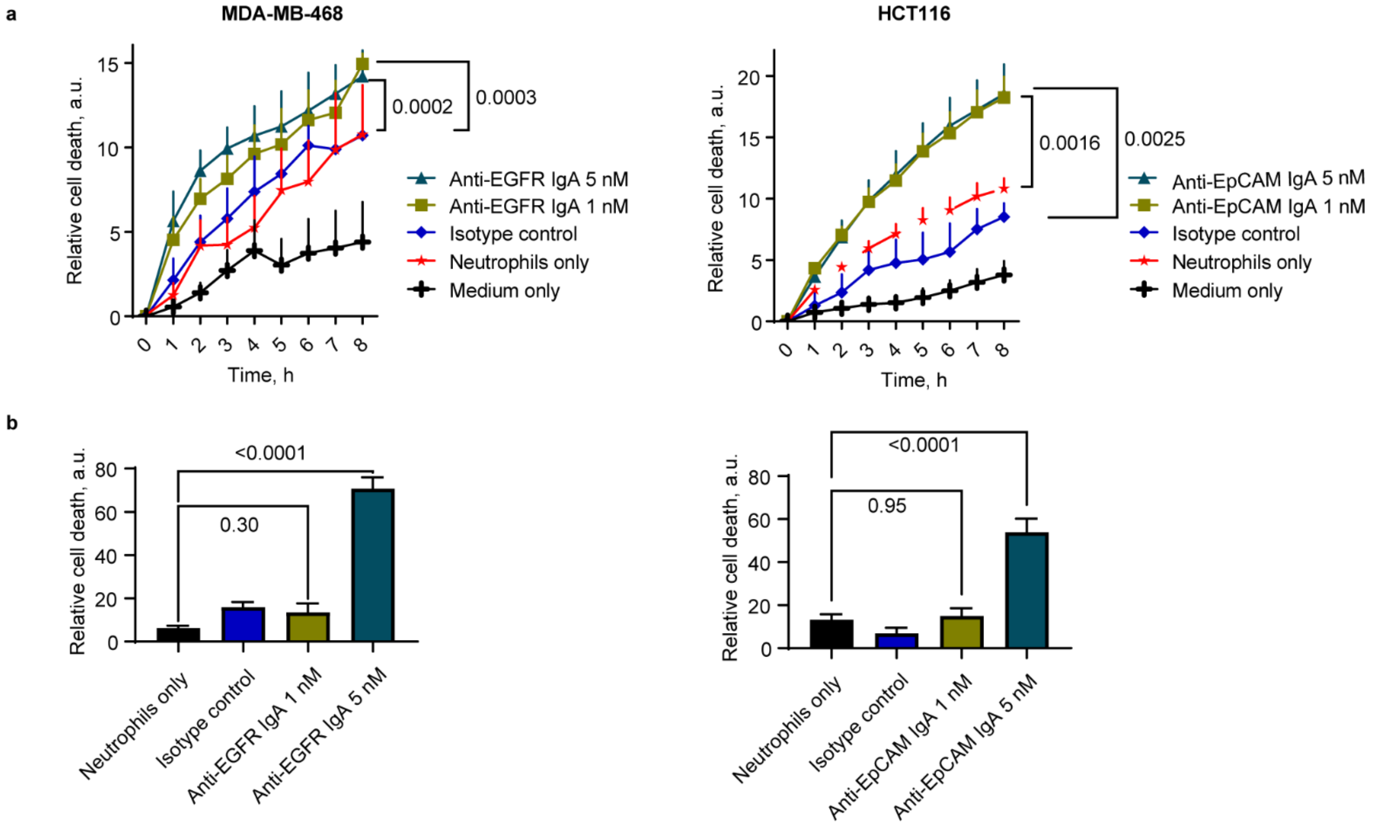


**Supplementary Figure S10. IgA antibody-activated primary human neutrophils kill cancer cells in monolayer and tumor-on-a-chip models. (a)** Time course of image-based ADCC analysis in monolayer models mediated via anti-EGFR IgA in MDA-MB-468, and via anti-EpCAM IgA in HCT116 cells. Recombinant IgA was used to assess the activity of the neutrophils co-cultured with tumor cells. As negative controls, a non-binding IgA (isotype control), or no antibody (neutrophils only) or no effector cells (medium only) were used. Each time point is shown as mean with SEM from four independent experiments, with effector cells from four different blood donors. Statistical analysis: one-way ANOVA with Dunnet's correction for multiple comparisons, adjusted p-values are shown. **(b)** ADCC in tumor-on-a-chip models (left, MDA-MB-468; right, HCT116). Anti-EGFR IgA and anti-EpCAM IgA were tested at the same concentrations as in (a), and neutrophils only (without antibody) or isotype control were used as negative controls. The values were normalized for the total sum per experiment and are shown as mean and SEM, for at least n=5 biological replicates (each value was obtained as an average from analyzing two representative chip areas, covering approx. 30% of the tissue compartment each), from three different blood donors. Statistical analysis: one-way ANOVA with Dunnet's correction for multiple comparisons, adjusted p-values are shown.

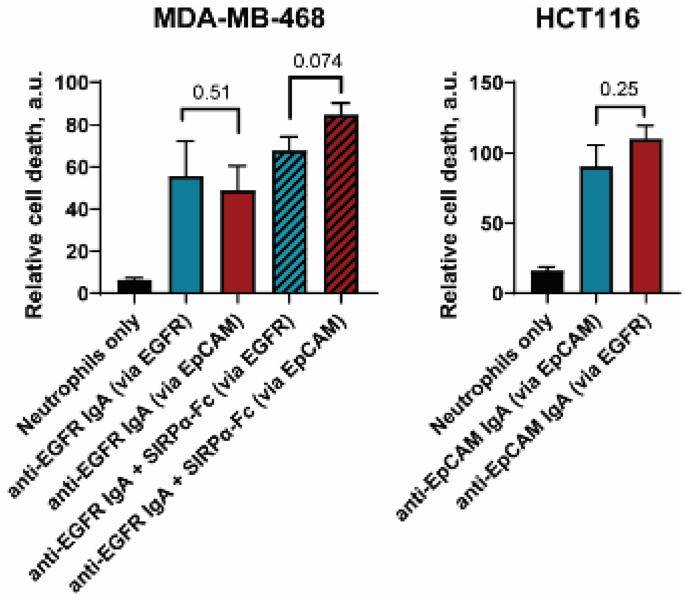


**Supplementary Figure S11. ADCC upon retargeting of adenoviral vector encoding IgA or SIRPα-Fc to cancer cells via EGFR or EpCAM in the tumor-on-a-chip.** ADCC activity was measured by an image-based assay (see Methods). In MDA-MB-468, ADCC was mediated via anti-EGFR IgA, and in HCT116, ADCC was mediated via anti-EpCAM IgA. As both EpCAM and EGFR could be targeted with adenovirus for the IgA (and SIRPα-Fc) gene delivery, both retargeting strategies were compared. Neutrophils without antibody served as a control. The values were normalized for the total sum of the experiment and are shown as mean and SEM, for at least n=5 biological replicates (each value was obtained as an average from analyzing two representative chip areas, covering approx. 30% of the tissue compartment each), from three different blood donors. Statistical analysis: unpaired t-test, adjusted p-values are shown.

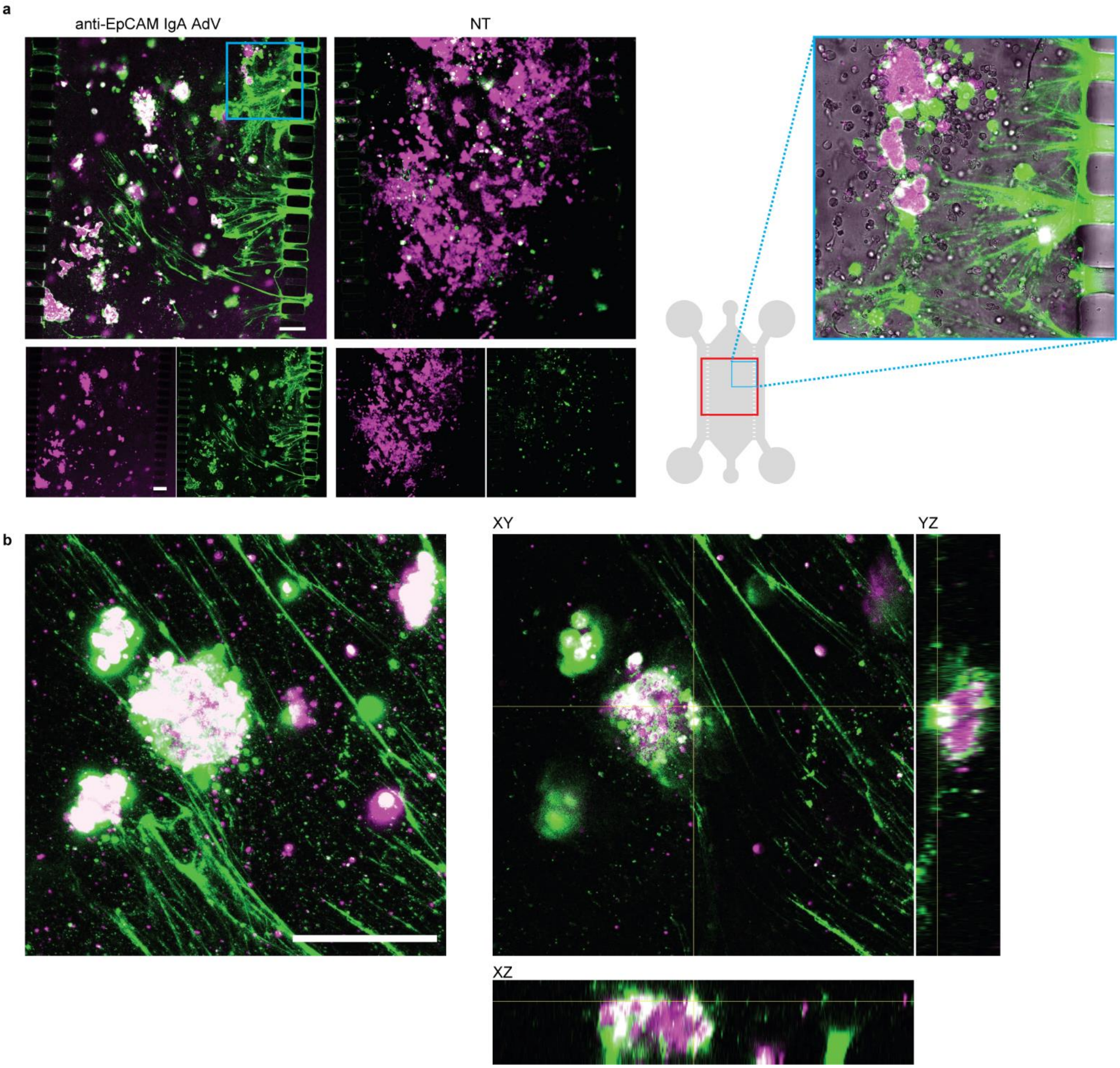


**Figure S12. Neutrophil extracellular traps (NETs) were detected on-chip following AdV-mediated local IgA production triggering neutrophil ADCC. (a)** Neutrophil activity is seen on-chip where anti-EpCAM IgA was delivered in the HCT116 model (left), versus non-treated (NT) control. Magenta: HCT116 cancer cells (CellTrace Far Red), green: DNA, non-viable cells (CellTox Green). Overlay images are shown in which co-localization of both signals indicate dead cancer cells; cells with only green signal represent non-viable neutrophils. Filamentous structures in the adenovirus-treated sample are likely neutrophil extracellular traps (NETs), which consist mostly of DNA, released by neutrophils which were perfused on-chip through the right-side channel. The schematic image represents the tumor-on-a-chip (not drawn to scale), with the approximate areas in the microscopy images shown in red; inset: a close-up of the tissue compartment showing the tunnel connections through which neutrophils infiltrate from the side channel. **(b)** A 3D reconstruction of cancer cells being eliminated by neutrophil ADCC, represented as a maximum intensity projection (left) and orthogonal sections (right). Representative images are shown, n=3. Scale bar: 100 µm.

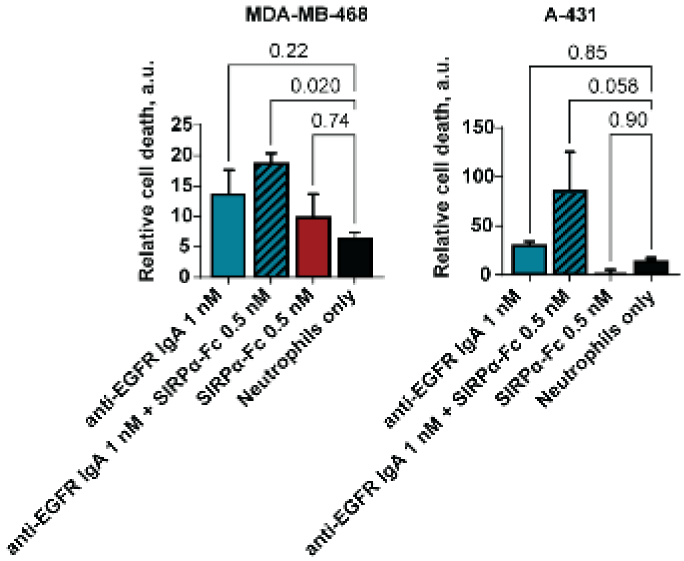


**Supplementary Figure S13. Effect of SIRPα-Fc protein on recombinant IgA-mediated neutrophil ADCC on-chip.** The effects of CD47 blockade were tested in MDA-MB-468 and A-431 cell lines, alone or in combination with anti-EGFR IgA. Neutrophils only: neutrophils added to the cancer cells without treatment. The values were normalized for the total sum of relative cell death values per experiment and are shown as mean ± SEM, for n=5 chips (MDA-MB-468), and n=2 chips (A-431). Each value was obtained as an average from analyzing two representative chip areas, covering approx. 30% of the tissue compartment each. Statistical analysis: one-way ANOVA with Dunnet's correction for multiple comparisons, adjusted p-values shown.

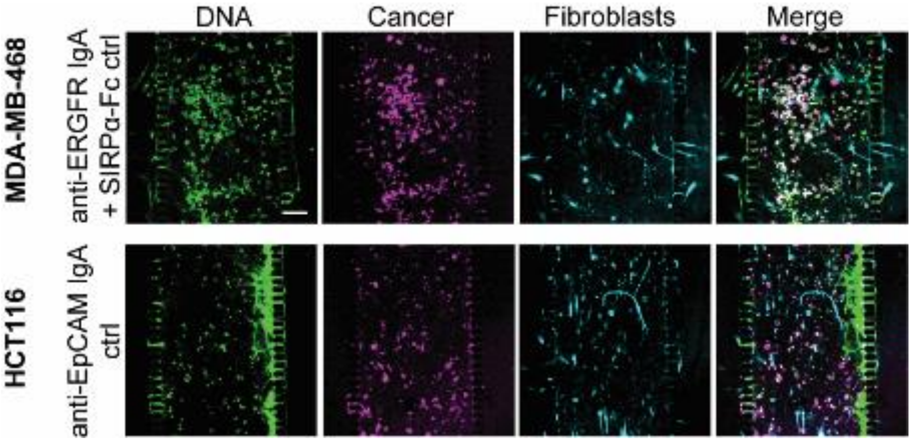


**Supplementary Figure S14. Specificity of IgA-based treatment in ADCC on-chip** (supplementary panel for Figure 6). Recombinant anti-EpCAM IgA (5 nM) or anti-EGFR IgA with SIRPα-Fc protein (1 nM and 0.5 nM, respectively) were added to HCT116 or MDA-MB-468 co-cultures with primary human skin fibroblasts on-chip. Labeled cancer cells (CellTrace Violet, magenta) were co-cultured on-chip in collagen with labeled healthy fibroblasts (CellTrace Far Red, cyan). The cell death was detected using DNA-binding CellTox Green dye (green). The overlay images demonstrate specificity for cancer cell killing compared to fibroblasts, since the CellTox stain overlaps with the magenta stain of the cancer cells. Scale bar: 200 μm. Representative images are shown from two chips from two independent experiments.

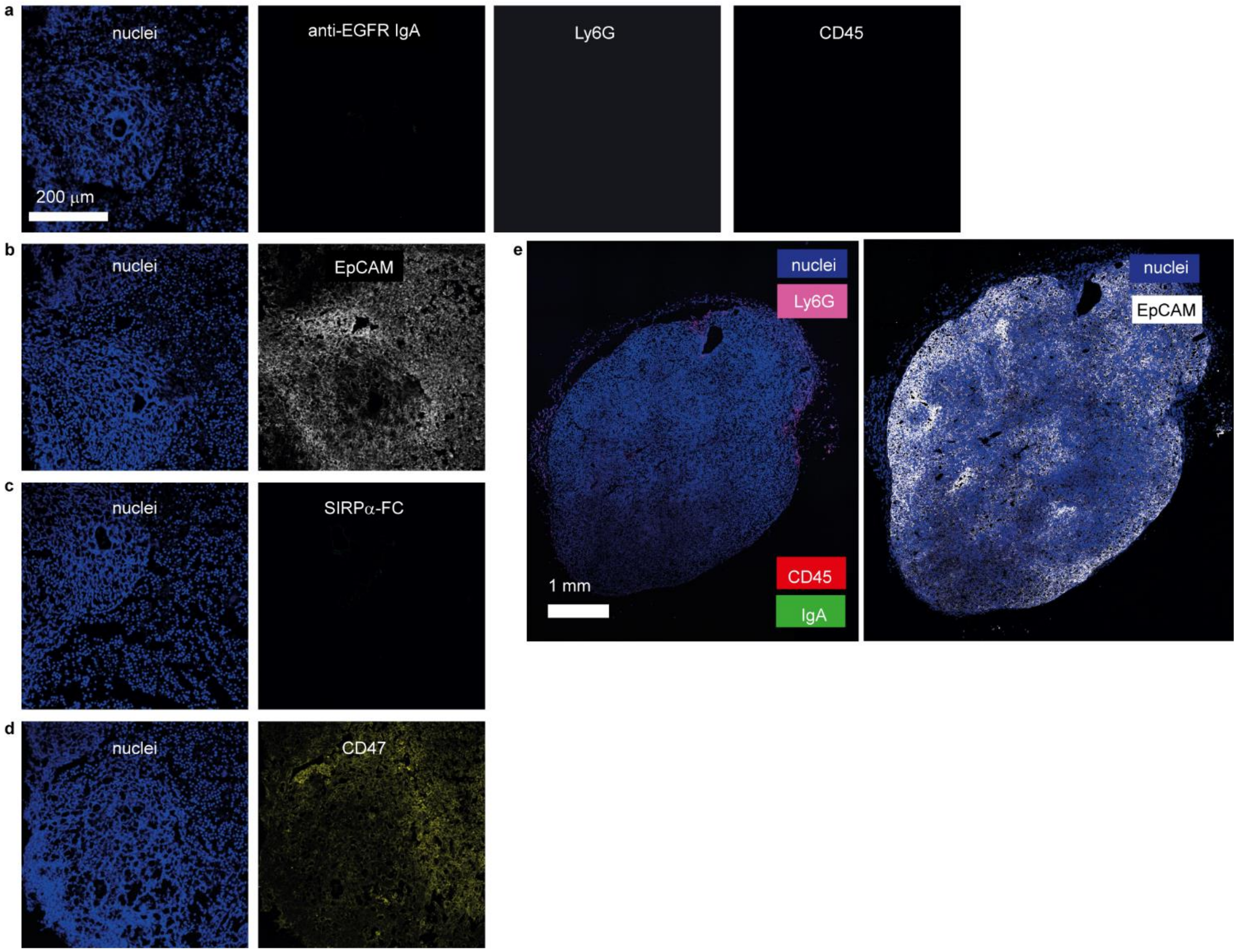


**Supplementary Figure S15. Immunofluorescence analysis of MDA-MB-468 tumor slices from SCID mice injected with PBS.** (supplementary panels for Figure 9)**. (a)** Immunofluorescence (IF) of tumor slices from MDA-MB-468 tumor-bearing SCID mice four days after injection with PBS. Nuclei were stained with DAPI. Ly6G and CD45 staining was detected in the same slice indicating the presence of neutrophils and immune cells, respectively. **(b-d)** Detection of EpCAM **(b)**, SIRPα-Fc **(c)** and CD47 **(d)** by IF as in (a). **(e)** A tile scan of a section of the entire tumor slice was analyzed by IF with the conditions described in (a) (left) or stained with anti-EpCAM IgA (right). The left image shows very little immune cell infiltration. The scale bar in (a) is applicable to all images from a-d. n = 4, representative images are shown.

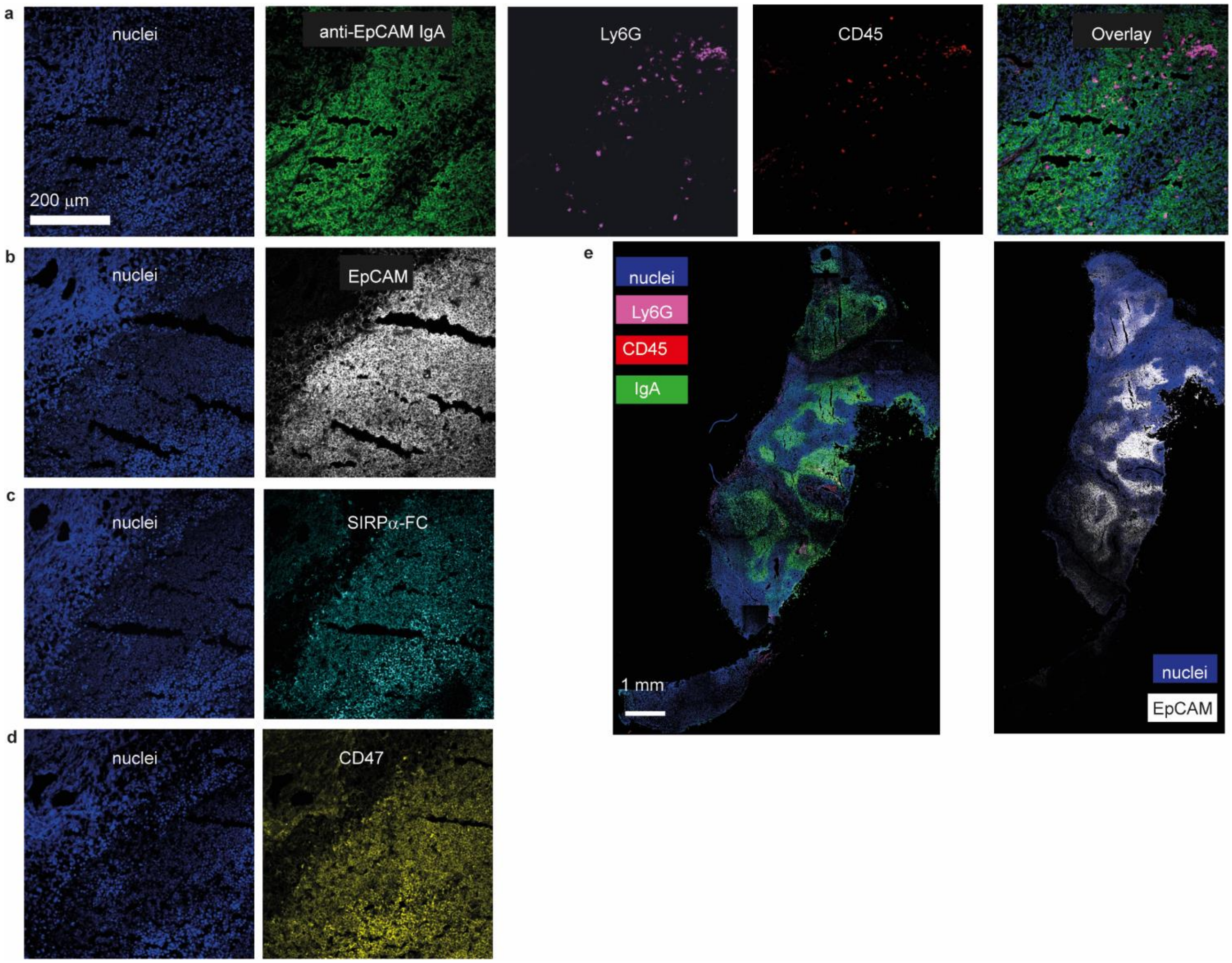


**Supplementary Figure S16. Immunofluorescence analysis of HCT116 tumor slices from SCID mice injected with anti-EpCAM IgA- and SIRPα-Fc-encoding adenoviral vectors** (supplementary panels for Figure 9)**. (a)** Immunofluorescence (IF) of tumor slices from HCT116 tumor-bearing SCID mice four days after injection with EGFR-directed anti-EpCAM IgA-encoding and SIRPα-Fc-encoding adenoviral vectors. Nuclei were stained with DAPI. Ly6G and CD45 were detected in the same slice as markers for neutrophils and immune cells, respectively. **(b-d)** Detection of EpCAM **(b)**, SIRPα-Fc **(c)** and CD47 **(d)** by IF as in (a). **(e)** A tile scan of a section of the entire tumor slice was analyzed by IF for the conditions described in (a) (left), or anti-EGFR IgA (right). The scale bar in (a) is applicable to all images from a-d. n = 6, representative images are shown.

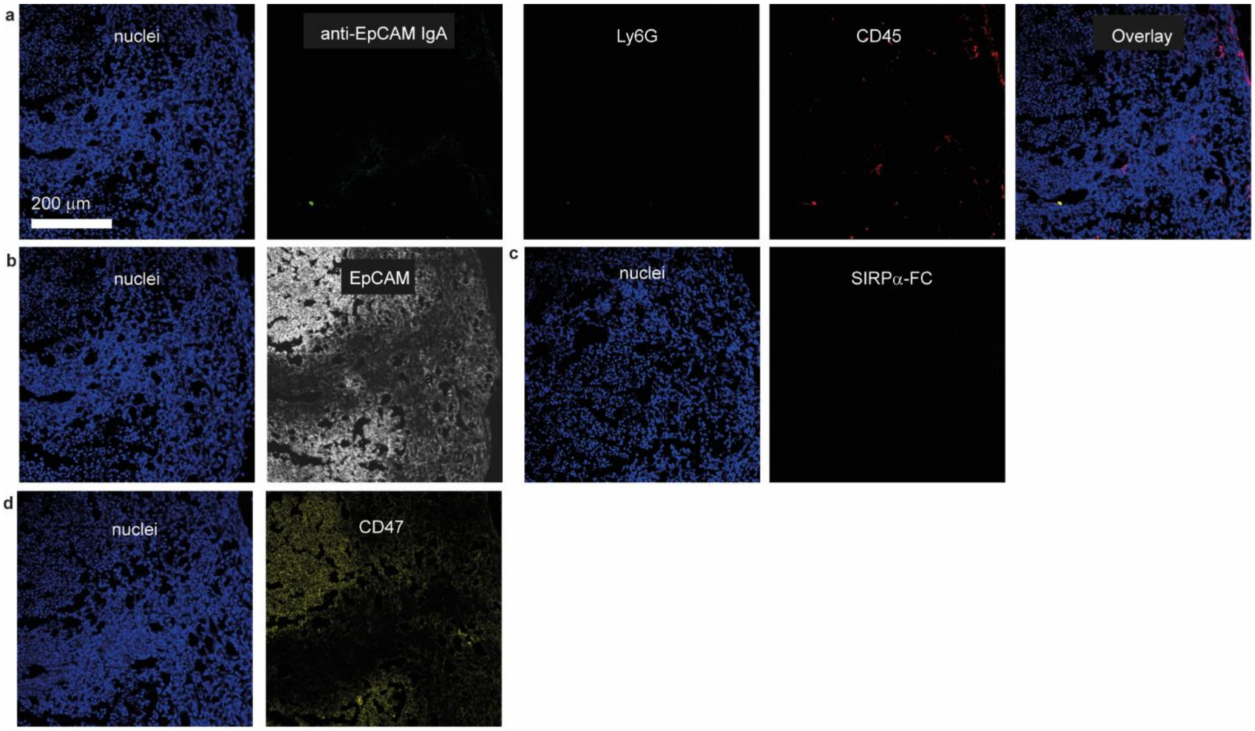


**Supplementary Figure S17. Immunofluorescence analysis of HCT116 tumor slices from SCID mice injected with PBS as control** (supplementary panels for Figure 9)**. (a)** Immunofluorescence (IF) of tumor slices from HCT116 tumor-bearing SCID mice four days after injection with PBS. Nuclei were stained with DAPI. Ly6G and CD45 were detected in the same slice as markers for neutrophils and immune cells, respectively. **(b-d)** Detection of EpCAM **(b)**, SIRPα-Fc **(c)** and CD47 **(d)** by IF as in (a). The scale bar in (a) is applicable to all images. n = 6, representative images are shown.

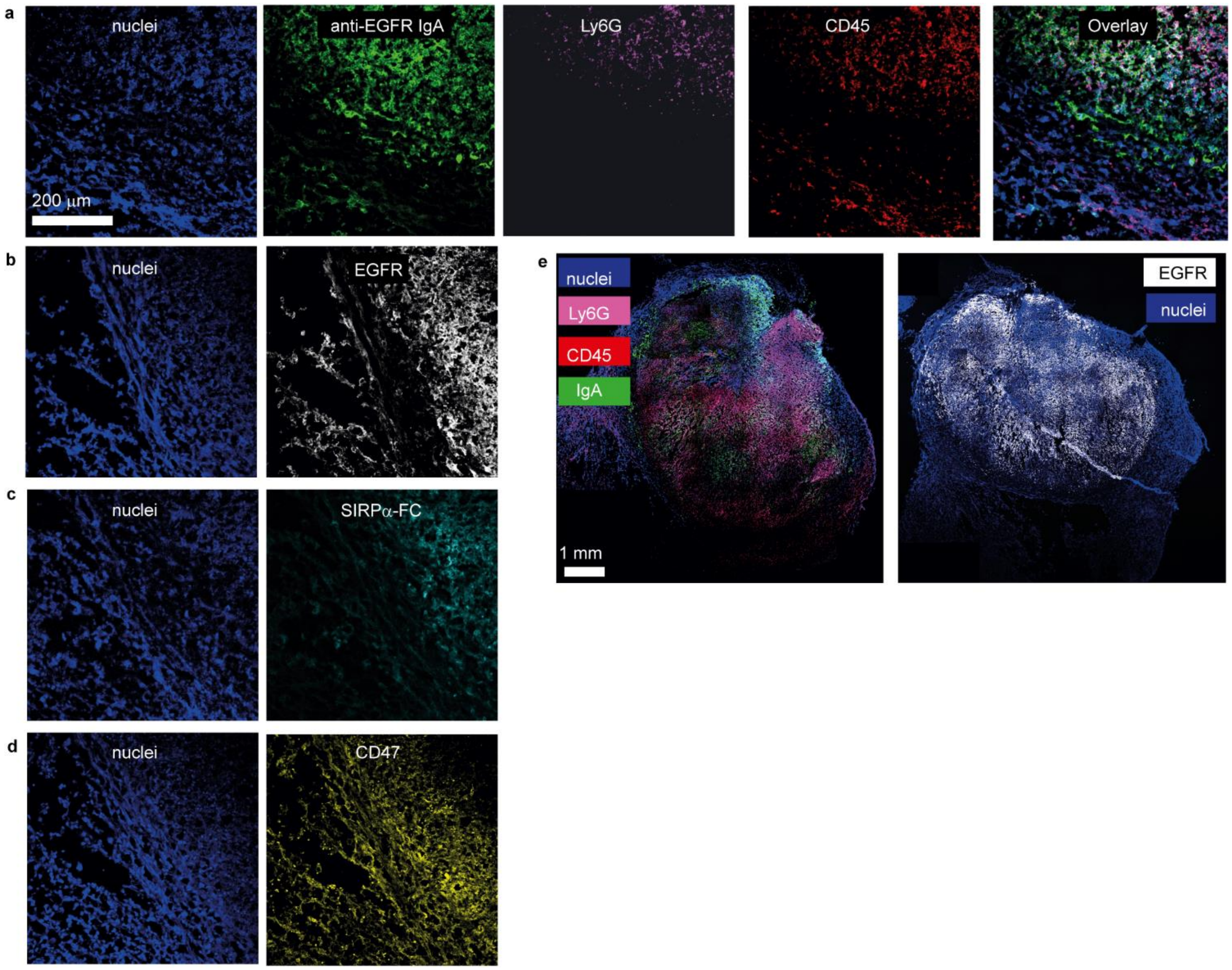


**Supplementary Figure S18. Immunofluorescence analysis of B16F10 tumors from C57BL/6 mice injected with anti-EGFR IgA- and SIRPα-Fc-encoding adenoviral vectors** (supplementary panels for Figure 9). **(a)** Immunofluorescence (IF) of tumor slices from B16F10 tumor-bearing C57Bl/6 mice four days after injection with EGFR-directed anti-EGFR IgA-encoding and SIRPα-Fc-encoding adenoviral vectors. Nuclei were stained with DAPI. Ly6G and CD45 were detected in the same slice as markers for neutrophils and immune cells, respectively. **(b-d)** Detection of EGFR **(b)**, SIRPα-Fc **(c)** and CD47 **(d)** by IF as in (a). **(e)** A tile scan of a section of the entire tumor slice was analyzed by IF for the condition described in (a) (left), or EGFR (right). The scale bar in (a) is applicable to all images from a-d. n = 6, representative images are shown.

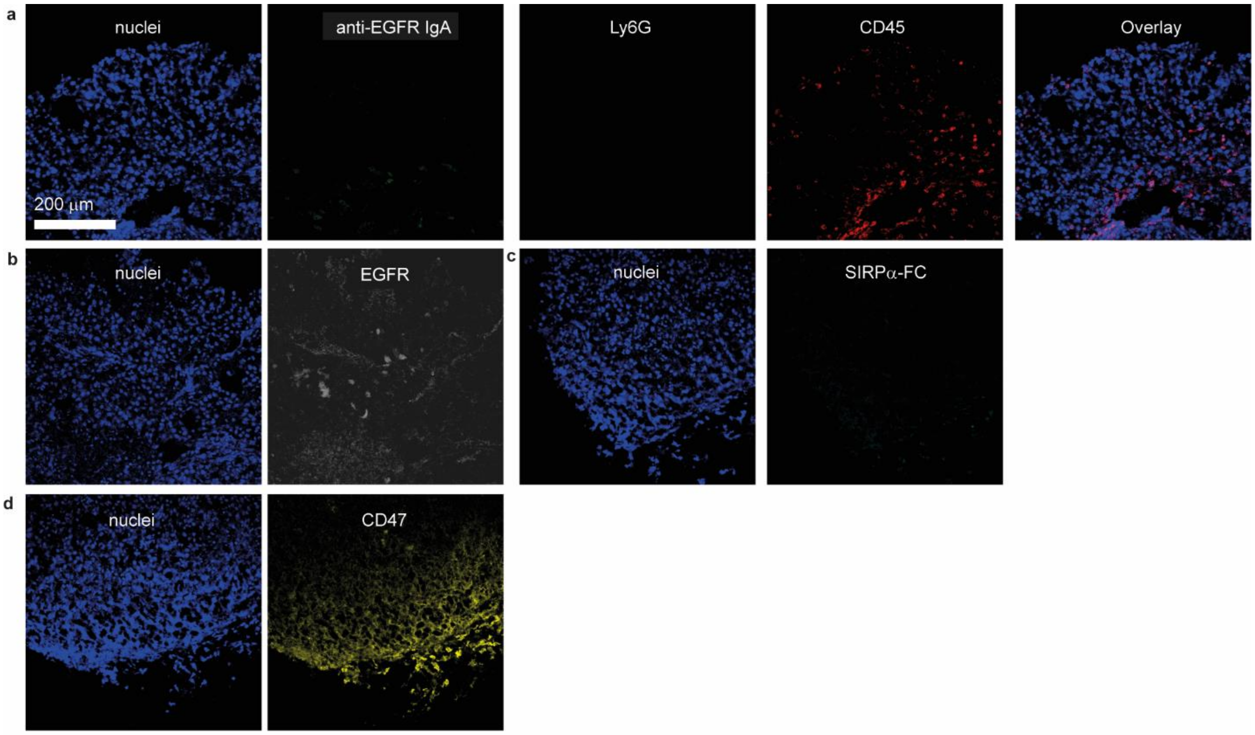


**Supplementary Figure S19. Immunofluorescence analysis of B16F10 tumors from C57BL/6 mice injected with PBS as control** (supplementary panels for Figure 9). **(a)** Immunofluorescence (IF) of tumor slices from B16F10 tumor-bearing C57Bl/6 mice four days after injection with PBS. Nuclei were stained with DAPI. Ly6G and CD45 were detected in the same slice as markers for neutrophils and immune cells, respectively. **(b-d)** Detection of EGFR **(b)**, SIRPα-Fc **(c)** and CD47 **(d)** by IF as in (a). The scale bar in (a) is applicable to all images. n = 6, representative images are shown.

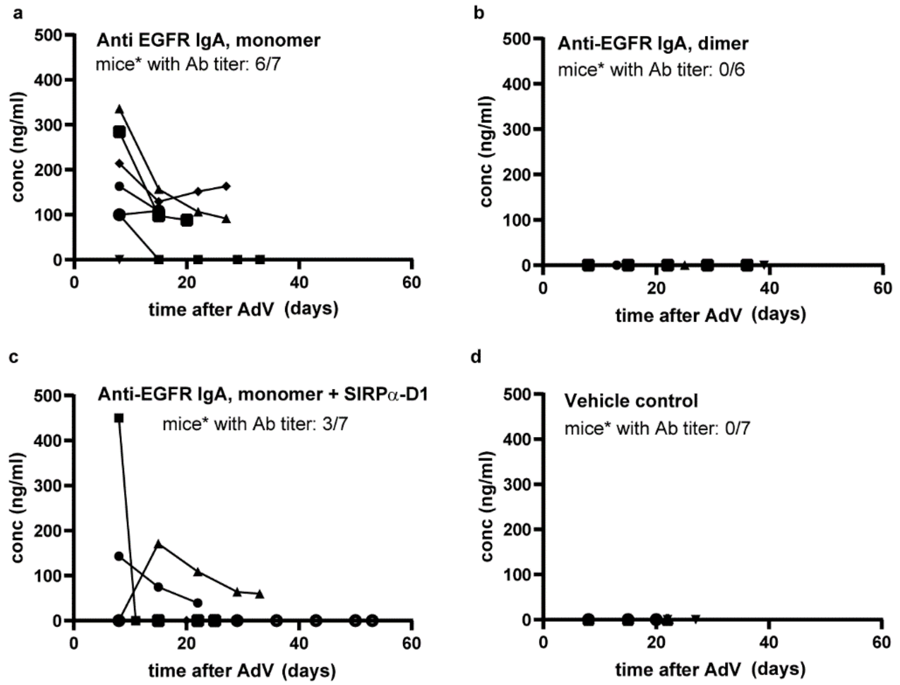


**Supplementary Figure S20. IgA antibody titers from MDA-MB 468 tumor-bearing SCID FcαRI transgenic mice injected with anti-EGFR IgA- and SIRPα-Fc-encoding adenoviral vectors.** IgA titers in plasma samples from tumor-bearing mice were measured by ELISA using monomeric IgA as a standard, with anti-human kappa light chain used for coating and anti-human IgA-HRP used for detection. Anti-EGFR IgA levels were detected upon treatment with (**a**) monomeric anti-EGFR IgA-encoding adenoviral vectors, (**b**) dimeric anti-EGFR IgA encoding adenoviral vectors, (**c**) monomeric anti-EGFR IgA and SIRPα-Fc-encoding adenoviral vectors or (**d**) vehicle control. Each symbol represents a different mouse.

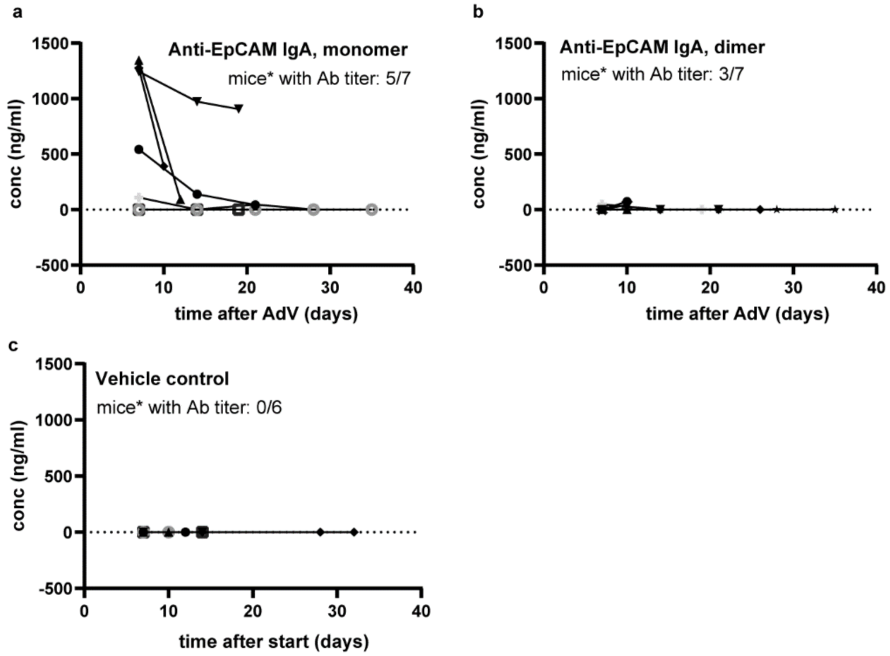


**Supplementary Figure S21. IgA titers from HCT116 tumor-bearing SCID FcαRI transgenic mice injected with anti-EpCAM IgA- and SIRPα-Fc-encoding adenoviral vectors.** IgA titers in plasma samples from tumor-bearing mice were measured using ELISA using monomeric IgA as a standard, with anti-human kappa light chain used for coating and anti-human IgA-HRP used for detection. Anti-EpCAM IgA levels were detected upon treatment with (**a**) monomeric anti-EpCAM IgA-encoding adenoviral vectors, (**b**) dimeric anti-EpCAM IgA encoding adenoviral vectors or (**c**) vehicle control. Each symbol represents a different mouse.

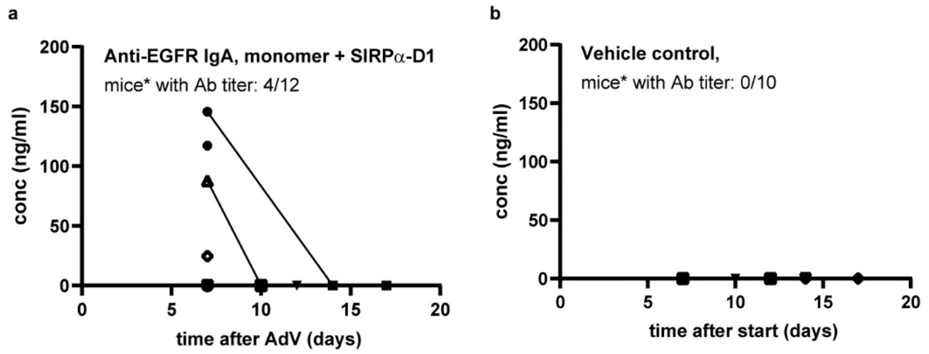


**Supplementary Figure S22. IgA titers from B16F10 tumor-bearing C57BL/6 FcαRI transgenic mice injected with anti-EGFR IgA- and SIRPα-Fc-encoding adenoviral vectors.** IgA titers in plasma samples from tumor-bearing mice were measured using ELISA using monomeric IgA as a standard, with anti-human kappa light chain used for coating and anti-human IgA-HRP used for detection. Anti-EGFR IgA levels were detected upon treatment with (**a**) monomeric anti-EGFR IgA and SIRPα-Fc-encoding adenoviral vectors or (**b**) vehicle control. Each symbol represents a different mouse.

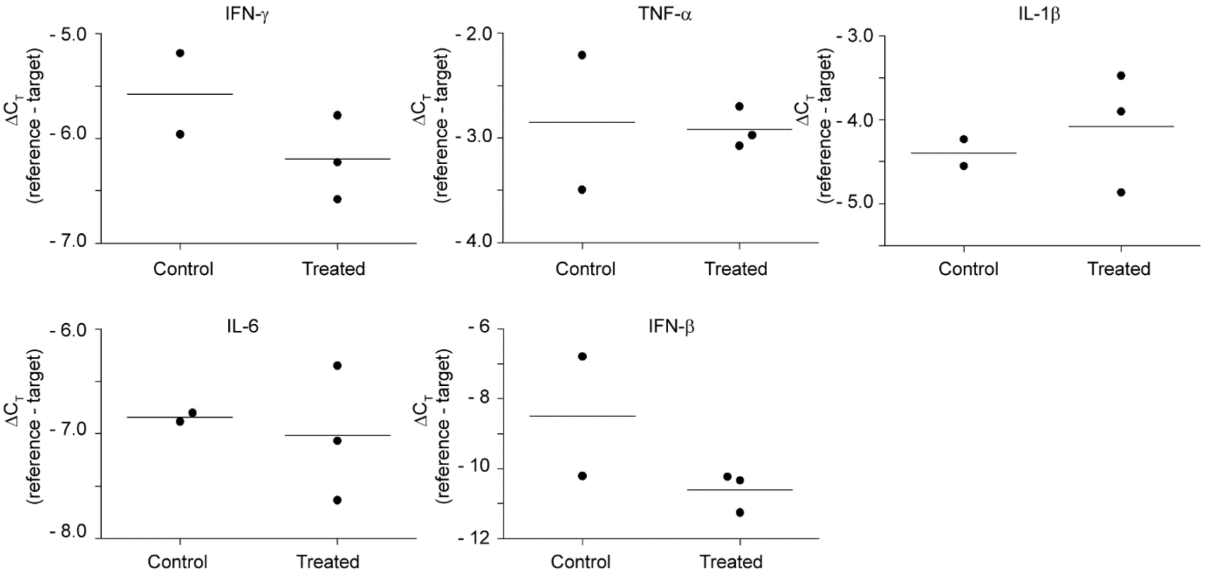


**Supplementary Figure S23. Gene expression analysis of B16F10 tumors extracted from C57BL/6 FcαRI transgenic mice.** Gene expression of indicated immune-associated markers was determined using qPCR from tumors extracted at day 32 from mice that were vehicle treated or that were injected with anti-EGFR IgA- and SIRPα-Fc-encoding adenoviral vectors. Each dot represents a different mouse.
